# A multireceptor six-phage cocktail consistently controls bacterial leaf spot of lettuce caused by *Xanthomonas hortorum* pv. *vitians* and improves harvest quality

**DOI:** 10.64898/2026.08.19.745710

**Authors:** Anaelle Baud, Inès Rougis, Danis Abrouk, Hanane Amari, Clara Aubremaire, Denis Costechareyre, Marie Graindorge Beaume, Alexandre Burlet, Franck Bertolla

**Author notes:** Correspondence: Franck Bertolla,; Anaelle Baud. Inès Rougis, Danis Abrouk, Hanane Amari, Clara Aubremaire, Denis Costechareyre, Marie Graindorge Beaume, Alexandre Burlet.

## Abstract

Phage cocktails are promising biocontrol agents against bacterial plant diseases by broadening host range and limiting the emergence of resistant mutants. To date, nine lytic phages with properties suitable for biocontrol have been isolated against *Xanthomonas hortorum* pv. *vitians*, the causal agent of bacterial leaf spot of lettuce. Here, a six-phage cocktail was rationally designed based on complementary host ranges, covering 91% of tested *vitians* strains while maintaining strict phage specificity toward the pathovar. To design a robust biocontrol, three distinct phage infection strategies, identified by transposon insertion sequencing, were combined in a cocktail. The susceptibility determinants were involved in LPS biosynthesis, a modified O-antigen structure, and an outer membrane protein putatively linked to the type I secretion system. As these structures contribute to plant colonization and virulence, phage resistance is expected to impose substantial fitness costs. In growth-chamber experiments, the phage cocktail provided dose-dependent protection, with significant symptom reduction observed across all tested concentrations, from 17% at 10^6^ PFU.mL^-1^, to 34.7% at 10^7^ PFU.mL^-1^ (two applications), and up to 66% at 10^8^ PFU.mL^-1^. In two independent field trials conducted across contrasting growing seasons, weekly applications consistently reduced disease severity by 30%, decreased the proportion of non-marketable lettuce heads by more than 84%, and reduced post-harvest trimming losses from 20.7% to 18.1% in summer and from 17.8% to 14.0% in autumn. These findings provide the first demonstration of a reproducible and effective phage-based biocontrol strategy against *Xanthomonas hortorum* pv*. vitians* under field conditions.

**IMPORTANCE:** In the context of growing food security challenges, sustainable strategies to control bacterial plant diseases are increasingly needed, particularly against *Xanthomonas* species, a leading genus of bacterial plant pathogens. Among these is *Xanthomonas hortorum* pv. *vitians*, the causal agent of bacterial leaf spot of lettuce, which occurs worldwide and for which no effective disease management strategies currently exists. While bacteriophages represent promising biocontrol agents, reliable phage-based treatments require overcoming major limitations, including narrow host range, bacterial resistance, and poor persistence under field conditions. Here, a six-phage cocktail combining complementary host ranges and distinct infection strategies was rationally designed and incorporated into a formulation containing an adjuvant already used in agriculture. The formulated phage cocktail provided disease protection under controlled conditions and reproducible efficacy across two field trials conducted in contrasting growing seasons. This study provides a framework for translating rational phage cocktail design into practical crop protection.

## INTRODUCTION

In the context of a growing global production, reducing crop losses caused by bacterial plant pathogens is an important component of sustainable agricultural production. Among affected crops, lettuce (*Lactuca sativa*, Asteraceae) is one of the most widely cultivated and consumed leafy vegetables worldwide (2), with a global production of about 28.2 million tons in 2024 and an estimated gross production value of 19.2 billion USD, including chicory (FAOSTAT crop database, https://www.fao.org/faostat/en/#data). The main producers are China, the United States, India, Italy, and Spain. However, lettuce production is threatened by several bacterial and fungal phytopathogens (3). Among the main bacterial pathogens, *Xanthomonas hortorum* pv. *vitians*, the causal agent of bacterial leaf spot of lettuce, was first reported in South Carolina in 1918 (4) and is now present on all continents where lettuce is cultivated (5). Under temperate and humid conditions (6, 7), the disease manifests as scattered, water-soaked lesions on lettuce leaves, which darken to brown or black, enlarge, and frequently coalesce (8). These symptoms can reduce head size and quality, ultimately leading to post-harvest losses that may reach 100% during major outbreaks (8, 9). The pathogen enters the plant through stomata, hydathodes, or wounds and can remain viable for several months on weeds, irrigation water, or infected plant debris on the soil surface (7, 10). Currently, no effective chemical control method is available against bacterial leaf spot of lettuce. Conventional compounds, such as copper-based products, require application at phytotoxic concentrations to be effective and raise environmental concerns, limiting their use in sustainable agriculture (11, 12).

As a result, novel control strategies that are environmentally sustainable and suitable for managing outbreaks of bacterial leaf spot of lettuce are needed. One promising alternative involves bacteriophages, viruses that selectively infect and lyse bacteria. Despite being discovered over a century ago and recognized for their biocontrol potential (13), bacteriophages have only recently regained attention in plant disease management (14). Phages possess several advantageous features for biocontrol applications, including their ability to self-replicate in the presence of the host, high host specificity, lack of toxicity to eukaryotic cells or production of toxic by-products, and overall ease of production (15). While their high host specificity minimizes off-target effects, it also represents a limitation for broad application. Bacterial susceptibility can vary substantially due to genetic diversity, particularly polymorphism in phage receptors, meaning that a phage may fail to infect all strains within a given species (16). In applications, naturally occurring phages are generally preferred over genetically engineered broad host range phages because of regulatory approval pathways and consumer acceptance (17). Furthermore, the antagonist nature of phage-bacteria interactions has led to an evolutionary arms race, with bacteria developing diverse resistance mechanisms (e.g., receptor modification or loss, production of extracellular matrix barriers, and adaptive immune systems) that can interfere with multiple steps of the lytic cycle (18). The development of phage cocktails contributes to addressing both challenges by broadening the host range and, through the combination of phages targeting distinct receptors or infection pathways, limiting the emergence and fixation of resistance mutations within bacterial populations. Ideally, a cocktail should cover the widest feasible set of target strains and combine phages with distinct receptor usage and infection mechanisms, ensuring complementary infectivity on the pathogen (19). Several strategies have been proposed to formulate robust phage cocktails, including stepwise screening and combination of candidates (20), selection of phages that target distinct bacterial receptors (21), incorporation of host-range mutant (h-mutant) phages (22, 23), and selection based on lytic activity profiles (24). Numerous phage cocktails have demonstrated protective efficacity against plant bacterial diseases under both greenhouse and field conditions. For instance, a four-phage cocktail targeting *Ralstonia solanacearum* reduced bacterial wilt incidence in tomato by up to 80% under greenhouse and field experiments. Disease suppression improved significantly with increasing phage richness: the four-phage cocktail provided the greatest reduction in disease incidence, outperforming combinations of two or three phages, as well as single-phage treatments (25). Similarly, promising results have been reported for cocktails targeting various *Xanthomonas* pathovars. For example, a three-phage cocktail targeting *Xanthomonas axonopodis* pv. *alii* reduced disease symptoms by 21.6-28.4% in field trials (26). Likewise, for *Xanthomonas arboricola* pv. *juglandis*, a high-dose treatment with a three-phage cocktail showed comparable efficacy to copper sulfate under field conditions (27). Recently, the first bacterial receptors required for infection by the phage Phi*Xhv*-1 infecting *X*. *hortorum* pv. *vitians* were identified using transposon insertion sequencing (Tn-seq), and were also shown to be involved in the bacterium’s infectivity on plants (28). In addition, eight new phages were characterized and revealed promising biocontrol-related properties *in vitro*. Notably, one of these phages was isolated on a mutant resistant to Phi*Xhv*-1 and specifically recognizes a mutated LPS structure (29).

In this study, we rationally designed a six-phage cocktail specifically targeting *X*. *hortorum* pv. *vitians* by combining phages with complementary host ranges and previously characterized genomic and functional characteristics. To elucidate the molecular basis of host susceptibility and support cocktail design, we generated two high-density transposon mutant libraries in susceptible host strains and used Tn-seq to identify bacterial genes required for infection by four phages. This genome-wide analysis revealed both shared and phage-specific host determinants, providing mechanistic insight into receptor diversity and the potential robustness of the phage cocktail against resistance evolution. The efficacy of the cocktail was subsequently evaluated through *in vitro* host-range assays and *in planta* experiments. Under controlled growth-chamber conditions, we assessed protection against bacterial leaf spot of lettuce across a range of phage concentrations and application frequencies. Finally, the biocontrol performance of the cocktail was validated in two independent field trials conducted across contrasting growing seasons, enabling assessment of both efficacy and reproducibility under agricultural conditions.

## MATERIALS AND METHODS

### Bacterial strains, plasmid and growth conditions

The *Escherichia coli* and *X*a*nthomon*as strains used in this study are listed in **Table S1**. *Xanthomonas* strains were routinely streaked onto tryptic soy agar (TSA) plates and incubated at 28°C for 48-72 h. Single colonies were used to inoculate tryptic soy broth (TSB) and cultured overnight at 28°C with shaking at 160 rpm. *E. coli* MFDpir/pSamEc was grown at 37°C in LB medium supplemented with diaminopimelic acid (DAP) at 57 µg.mL^-1^. The pSamEc plasmid is a mobilizable suicide vector encoding (i) an ampicillin resistance (*bla*), (ii) a Himar1-C9 transposase gene under the control of the lac promoter, and (iii) a kanamycin resistance gene flanked by mariner inverted repeat sequences containing *MmeI* restriction sites. When required, media were supplemented with 100 µg.mL^-1^ ampicillin, and 50 µg.mL^-1^ kanamycin. The strains were kept at −70°C in 30% (v/v) glycerol vials for long-term storage.

### Phage cocktail design and production

Based on their host range, lytic kinetics parameters, genomic characteristics, and abiotic stability, six previously characterized phages (i.e., Phi*Xhv*-1, Phi*Xhv*-2, Phi*Xhv*-5, Phi*Xhv*-7, Phi*Xhv*-12 and Phi*Xhv*-28) (28, 29) were selected to formulate a phage cocktail targeting *X. hortorum* pv. *vitians*. Each phage was amplified individually in liquid culture using its isolation host as previously described (28). Phage titers (PFU.mL^-1^) were determined using the double agar layer spot assay. Unless specified otherwise, the six phages were mixed at equal PFU concentrations immediately before use. The stability of each phage within the cocktail was verified by titration using selective bacterial hosts. In addition, a long-term storage assay was performed by keeping the phage cocktail in TSB at 4°C.

### *In vitro* growth inhibition assay of the phage cocktail and AUC-based quantification

The susceptibility of thirty-four *X. hortorum* pv. *vitians* strains and six strains belonging to other *Xanthomonas* pathovars or species to the phage cocktail was assessed using the Bioscreen C MBR system (Thermo Fisher Scientific), as previously described (30). Each condition was performed in five technical replicates. Raw absorbance values were corrected by subtracting the mean absorbance of TSB blanks at each time point. The area under the curve (AUC) was computed using the trapezoidal rule with custom R scripts. Bacterial growth inhibition was quantified as follows: Growth inhibition (%) = 100 × [(AUC_control_ – AUC_phage_)/AUC_control_], where AUC_control_ corresponds to bacteria grown without phages and AUC_phage_ to phage-treated cultures.

### Construction of transposon mutant libraries and genomic DNA extraction

To identify bacterial genes involved in phage infection and infer the corresponding receptors, three transposon mutant libraries were used, including a previously described library constructed in *X*. *hortorum* pv. *vitians* strain CFBP8638 (31). Two additional transposon mutant libraries were generated in strains CFBP8641 and CFBP8642 by biparental conjugation with the donor strain *E. coli* MFDpir/pSamEc carrying the Himar1-C9 mariner transposon, as described by Morinière *et al*. (31), with slight modifications. After centrifugation (3,344 x *g*, 10 min, 21°C), pellets of overnight cultures were washed with 0.8% NaCl. The centrifugation and washing steps were repeated three times. For strain CFBP8641, two conjugation mixtures were prepared using approximately 10^10^ cells of each donor and recipient cells, whereas fourteen conjugation mixtures were performed for strain CFBP8642 to compensate for its lower transposition efficiency. After centrifugation (3,344 x *g*, 20 min, 21°C), the pellets were resuspended in 250 µL TSB supplemented with DAP (57 µg.mL^-1^), and spotted onto over-dried TSA plates. After overnight incubation at 28°C, bacterial cells were recovered in TSB, plated onto TSA plates supplemented with kanamycin, and incubated for 72 h at 28°C to select transposon mutants. Colonies were harvested in TSB, pooled, mixed with sterile 30% (v/v) glycerol, and stored at −70°C. The resulting libraries contained an estimated 168,000 and 205,000 independent transposon mutants for strains CFBP8641 and CFBP8642, respectively, corresponding to final concentrations of 2.85 ×10^10^ and 3.61 × 10^10^ CFU.mL^-1^. Loss of the transposase gene following transposition was verified by PCR on a subset of mutants. Genomic DNA was extracted in duplicate from 125 µL aliquots of each library using the Wizard Genomic DNA Purification Kit (Promega).

### Genome-wide phage-resistance screening by Tn-seq

Transposon library aliquots were thawed at 4°C for 1 h prior to inoculation. Bacterial suspensions were adjusted to an OD_600nm_ of 0.15 (7.5 × 10^7^ CFU.mL^-1^) in 30 mL of TSB supplemented with 10 mM CaCl_2_, and incubated for 30 min at 28°C with shaking at 150 rpm. Each phage was then added to the corresponding transposon library (Phi*Xhv*-2 and Phi*Xhv*-5 to CFBP8642, Phi*Xhv*-7 to CFBP8641, and Phi*Xhv*-12 to CFBP8638) at a MOI of 1, or 5 for Phi*Xhv*-5. To minimize the enrichment of spontaneous phage-resistant mutants, wild-type cultures challenged with each phage were monitored in parallel to determine the maximum incubation time that maintained bacterial growth inhibition before resistant populations emerged. Accordingly, phage-treated transposon libraries were incubated at 28°C, 150 rpm, for 16 h (Phi*Xhv*-2 and Phi*Xhv*-5), or 24 h (Phi*Xhv*-7 and Phi*Xhv*-12), following a 30 min adsorption period. Control libraries were grown without phage. All conditions were performed in biological duplicates. Cell pellets were collected by centrifugation at 5,000 x *g* for 10 min at 4°C. Genomic DNA library extraction and preparation were conducted as described previously (31) and were submitted to the I2BC-sequencing platform (I2BC, Gif-sur-Yvette, France) to be sequenced in single-read 75-bp on a NextSeq 5000 instrument (Illumina, Inc.).

### Fitness assessment of genes associated with phage infection

Sequencing reads preprocessing and TnSeq analysis were performed as described previously (28). Briefly, read counts were normalized using the “Totreads” method implemented in TRANSIT v.3.3.19 (32) to obtain the same total number of reads for each sample. The contribution of each genetic feature to bacterial fitness during phage infection was determined by pairwise comparisons of phage-treated and control libraries using the “Resampling” method implemented in TRANSIT v.3.3.19. Reads mapping to the 5% N-terminal and 10% C-terminal portions of each genetic feature were discarded, and a LOESS (locally estimated scatterplot smoothing) correction was applied to account for genome positional bias. Genetic features with a log_2_ Fold Change (log_2_FC) > 2 and a *q-value* ≤ 0.05 were considered required for successful phage infection.

### Evaluation of phage cocktail efficiency against bacterial leaf spot of lettuce under growth chamber conditions

The efficacy of the phage cocktail against the bacterial leaf spot of lettuce was evaluated under growth chamber conditions on leaf lettuce cv. Météore. For each treatment condition, eight plants were used, and each experiment was independently repeated three times. Plants were grown in a greenhouse in 8-cm pots containing TS3 mold (Klasmann-Deilmann, Germany) for 3 weeks, until reaching the 5-6 true leaf stage. Each of the six phages was incorporated at equal concentrations into a formulation containing 1% (w/v) bentonite supplemented with 2% (v/v) Escapade® (ActionPin, Castets, France). For each application, 20 mL of phage cocktail suspension was applied to the aerial parts of plants using a hand sprayer. Three multiplicities of infection (MOI) were tested (0.01, 0.1, and 1) with either one or two applications. The phage cocktail was tested as a preventive treatment, applied 4 h prior to inoculation with the strain *X*. *hortorum* pv. *vitians* CFBP8638. A second phage application was performed 24 h after bacterial inoculation. Immediately after phage application, plants were transferred to a growth chamber (Ineltec, Spain) set at 25°C, in the dark and 90% relative humidity for 4 h. Plants were then inoculated by foliar spraying, with each treatment group receiving 40 mL of bacterial inoculum (around 5 x 10^7^ CFU/lettuce). The bacterial inoculum consisted of strain CFBP8638 grown overnight in one-tenth-strength TSB and adjusted to an OD_600nm_ of 0.2 (approximately 10^8^ CFU/mL) in sterile deionized water containing 0.08% Tween 80. For the infection control, plants received the formulation followed 4 h later by the bacterial inoculum as described above. The negative control received the formulation followed by the same inoculum solution without bacteria. Post-inoculation, plants were maintained in the growth chamber under a 12 h dark/12 h light photoperiod (12 290 lux) and 90% relative humidity (RH) for the first 48 h, followed by 70% RH until the end of the experiment. Disease severity was evaluated on each plant every 2-3 days, from 8 to 20 days post-inoculation (DPI), using a 0-5 symptom severity scale (33) (**Fig. S1**). For each MOI and treatment combination, disease severity scores from the eight plants within each independent biological replicate were averaged to obtain a replicate mean. For each day post-inoculation, the mean and 95% confidence interval across the three replicate means were calculated to generate disease progression curves.

### Field-based assessment of phage biocontrol efficiency

In 2025, two independent field trials were conducted at the CTIFL experimental station in Brindas, France (45°43’40.134’’N, 4°43’41.725’’E), one during the summer season and the other in early autumn. Weather conditions during both field trials are shown in **Figure S2**. The site is characterized by a sandy-loam-clay soil. The lettuce cultivar Météore was used. To establish endemic disease pressure, lettuce plants infected with *X*. *hortorum* pv. *vitians* strain CFBP8638 were crushed and incorporated into the soil two months before the start of the biocontrol assay. A second soil contamination was performed before transplanting. The main efficacy experiment comprised three treatments: (i) phage-treated plots (pre-inoculated and receiving the phage cocktail formulated in 1% bentonite and 2% Escapade®), (ii) an infected formulation control (pre-inoculated and receiving the same formulation without phage), and (iii) an uninfected formulation control (plots uninoculated, receiving the formulation without phage). To assess whether the formulation itself affected disease development or lettuce quality, two additional reference treatments were included, consisting of inoculated and non-inoculated plots that received no formulation. Each treatment consisted of four randomized plots (7.5 m^2^ each) containing 80 plants, totaling 320 plants per treatment (**Fig. S2.C**). Seedlings were transplanted at the 4-leaf stage. Ten days later, when plants reached the 7-leaf stage, the six-phage cocktail was applied using a Euro-pulvé wheeled sprayer (Aspach, France) equipped with a five-nozzle boom (Teejete XR 110015 VS, 25 cm spacing). The six-phage cocktail was applied at a final dose of 4 × 10^11^ PFU.m^2^. Hoeing was performed ten days after the first phage application. Phage applications were performed late in the day and repeated three times at weekly intervals. Whenever thunderstorms occurred (defined as rainfall > 5 mm and temperature > 25 °C), an additional application was performed immediately after the storm, after which the original weekly schedule was resumed. Disease severity was visually scored on twenty plants per elementary plot from the central row using a standardized five-point disease index (**Fig. S2.B**). At harvest, raw and trimmed head weights were recorded for 20 lettuce heads per plot.

### Statistical analyses

All statistical analyses were performed in R (v4.3.3; R Core Team) within RStudio (version 2023.12.1.402 “Ocean Storm” Release; RStudio Team).

For the growth-chamber experiment, disease severity at 20 dpi was analyzed separately for each MOI. Symptoms score were corrected by subtracting, within each replicate, the mean of the negative control before analysis. Corrected scores were analyzed using Gaussian linear mixed-effects models fitted with *glmmTMB*, including treatment as a fixed effect and replicate as a random intercept. Pairwise comparisons were obtained from estimated marginal means (*emmeans*) with Tukey adjustment. Protection (%) was calculated for each plant with the adjusted severity score and expressed relative to the corresponding infected control within each replicate and MOI. At 20 DPI, protection was analyzed using a linear model including MOI, number of phage applications, their interaction, and replicate as a blocking factor. Dunnett-type contrasts compared each phage treatment with the infected control within each MOI. Changes in protection over time (8 to 20 DPI) were analyzed using linear mixed-effects models (*nlme, REML*) including day, MOI, number of applications, and all interactions as fixed effects. Replicate and plant identity nested within replicate were included as random intercepts, and an AR(1) correlation structure accounted repeated measurements on the same plant. Sidak adjusted Dunnett contrasts compared each assessment date with the 8 DPI baseline.

For the field experiments, symptom severity (0-4) was analyzed using cumulative link mixed models (CLMM; *ordinal* package) treating symptom score as an ordinal response. Treatment was included as a fixed effect and block as a random intercept. The probability of producing unmarketable lettuce (score ≥ 3) was analyzed using binomial generalized linear mixed models (GLMM). Fresh and trimmed weights were analyzed using linear mixed-effects models (*lme4*), whereas trimming loss proportion was analyzed using beta generalized linear mixed models. For analyses assessing the effect of the formulation alone, inoculation status, formulation, season, and their interactions were included as fixed effects, with plot nested within season as a random intercept. Pairwise comparisons were obtained from *emmeans*, using Tukey adjustment for treatment comparisons and Holm correction when comparing formulated and non-formulated controls.

For analysis, model assumptions were assessed by visual inspection of resudial diagnostics. Core R packages included *glmmTMB* v1.1.11, *ordinal* v2023.12-4.1, *lme4 v*1.1-37, *nlme* v3.1-164, *emmeans* v1.11.2, and *multcomp* v1.4-28. Data import and preprocessing used *readxl* v1.4.5, *readr* v2.1.5, *stringr* v1.5.1, and *dplyr* v1.1.4. Figures were generated with *ggplot2* v3.5.2 and finalized using Inkscape v1.4.

## RESULTS

### Rationale, efficacy *in vitro*, and stability of the phage cocktail

Drawing on a previously characterized phage collection (29), a six-phage cocktail was assembled to maximize coverage of the genetic diversity of *X*. *hortorum* pv. *vitians* while maintaining high host specificity. The cocktail combined phages with complementary and partially overlapping host ranges, with Phi*Xhv*-28 specifically included to cover mutants resistant to Phi*Xhv*-1. Based on the previously reported individual host-range profiles, 31 of the 34 strains (91%) were susceptible to at least one phage. Among them, 8 strains (23%) were susceptible to two phages, 5 strains (15%) to three phages, and one strain to five phages. Notably, Phi*Xhv*-12, Phi*Xhv*-5, and Phi*Xhv*-28 uniquely extended the overall host range of the cocktail by infecting 13, 3, and 1 additional strains, respectively, that were not susceptible to any other phage in the cocktail (**Fig.1.A**). This combination was intended to close host-range gaps and limit the emergence of spontaneous resistance. In liquid culture assays, the host-range profile of the formulated cocktail matched the predicted combined host range of the six individual phages, indicating that no antagonistic interactions occurred between phages following formulation. Accordingly, the cocktail strongly inhibited 31 of the 34 *Xanthomonas hortorum* pv. *vitians* strains (> 70% inhibition), while the remaining three strains (i.e., LM17388, LM17422, and LM17692) were completely resistant, resulting in an overall coverage of 91% of the tested collection. No lytic activity was detected against six non-*vitians* strains (< 10% inhibition), confirming the strict specificity of the cocktail (**Fig.1.B**). Cocktail compatibility and stability were further supported by long-term storage experiments. After one year at 4°C, all six phages remained detectable on their respective indicator strains, with changes in titer ranging from −1.38 log_10_ PFU.mL^-1^ (Phi*Xhv*-2) to +0.22 log_10_ PFU.mL^-1^ (Phi*Xhv*-28) relative to their initial theoretical concentrations. The overall cocktail titer decreased by 0.25 log_10_ PFU.mL^-1^, indicating that the six phages remained compatible during storage, with no evidence of competitive exclusion or preferential loss of individual phages (**Table S2**).

**Figure 1.**
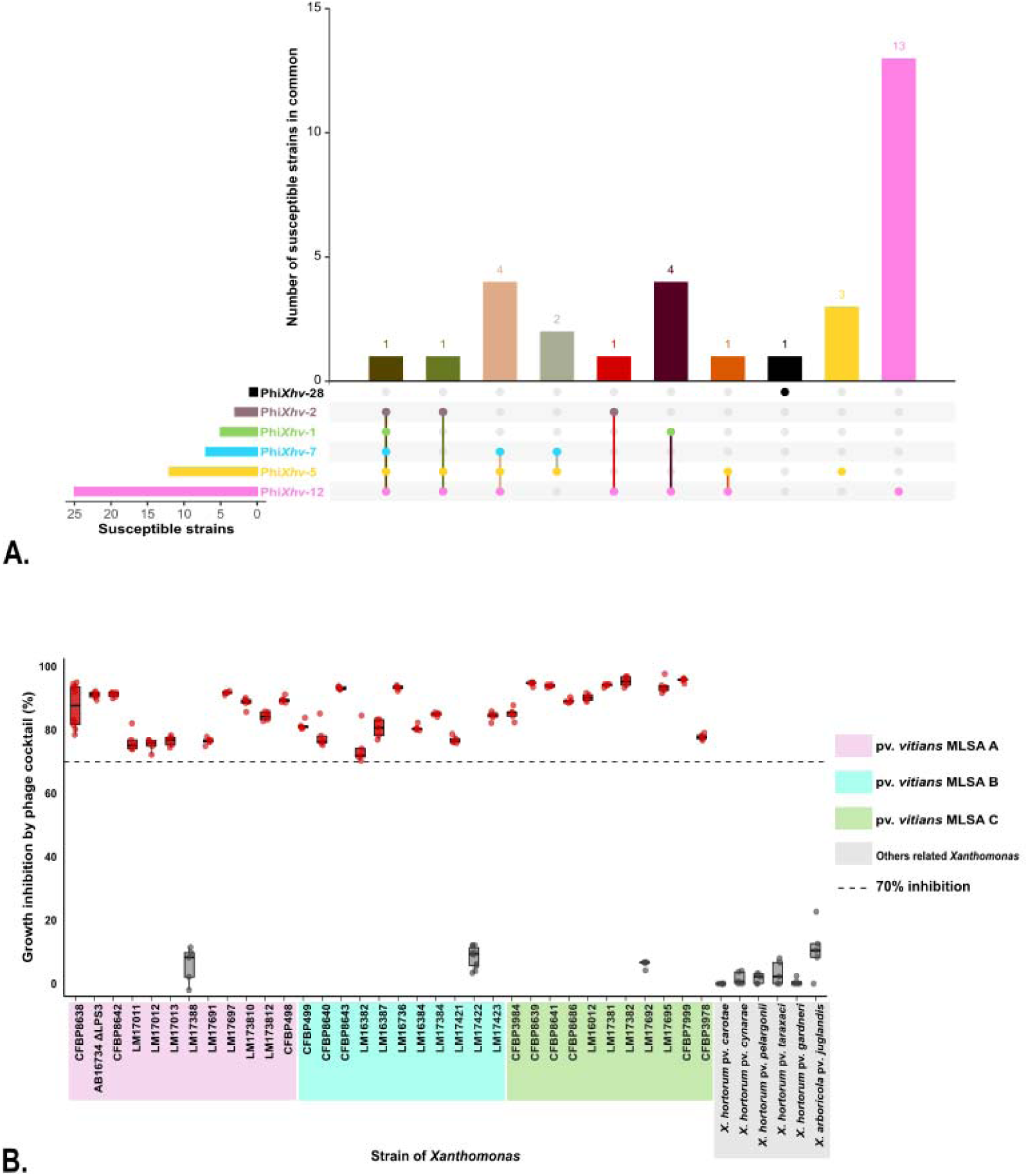
Rational design and *in vitro* validation of a six-phage cocktail targeting *X*. *hortorum* pv. *vitians*. **A.** UpSetplot illustrating the overlap in host range of six selected phages (Phi*Xhv*-1, Phi*Xhv*-2, Phi*Xhv*-5, Phi*Xhv*-7, Phi*Xhv*-12, and Phi*Xhv*-28), based on previously published spot assay data (29). Only strains with an efficiency of plating (EOP) > 10^-4^ were considered susceptible. Horizontal bars indicate the number of strains susceptible to each individual phage. Vertical bars indicate the number of susceptible strains corresponding to each host-range intersection, with colored dots and connecting lines identifying the phage or combination of phages defining each intersection. **B.** *In vitro* efficacy of the six-phage cocktail in liquid medium against a diverse panel of *Xanthomonas* strains. Growth inhibition was quantified as the percentage reduction in area under the growth curve (AUC) at 20 h relative to untreated controls. The grey dashed line marks the 70% inhibition threshold. Grey boxplots represent strains with <70% growth inhibition, whereas red boxplots represent strains with >70% growth inhibition. Individual replicates are shown as dots. Strains are grouped according to species and multilocus sequence analysis (MLSA) classification: pink = MLSA group A, cyan = MLSA group B, green = MLSA group C, and grey = other phylogenetically related *Xanthomonas* species or pathovars.

### Construction and quality assessment of Tn-seq mutant libraries

To identify the bacterial determinants underlying the complementary infection strategies of the cocktail phages, each phage was individually challenged against a Tn-seq library generated from a fully susceptible host strain. Receptor determinants for Phi*Xhv*-1 were available from a previous Tn-seq study (28), whereas Phi*Xhv*-28 was not investigated because it specifically infects mutants carrying alterations in the O-antigen rather than the corresponding wild type strain (29). Consequently, two new transposon mutant libraries of strains CFBP8641 and CFBP8642 were generated using the Himar1 transposon carried by plasmid pSAM_Ec (**Table S1**). On average, 87.3% and 78.0% of sequencing reads contained the Himar1 transposon end sequence, of which 99.6% and 99.3% mapped to unique TA sites in the CFBP8641 and CFBP8642 genomes, respectively (**Fig. S3.A**). The near-complete mapping of transposon-end-containing reads to unique TA sites indicated the absence of residual delivery vector sequences, which could otherwise have markedly reduced the number of informative reads. Mapping of filtered reads revealed that, out of the 83,244 and 79,689 TA dinucleotide target sites in the genomes of CFBP8641 and CFBP8642, an average of 58,480 and 51,004 TA sites carried at least one insertion, respectively (**Table S3**). This corresponds to mean insertion densities of 70.25% and 64%, i.e., approximately one insertion every 90 bp and 102 bp, respectively. Pearson correlation coefficients between biological replicates were high, 0.89 for CFBP8641 and 0.93 for CFBP8642, confirming the robustness and saturation of both libraries for genome-wide screening of phage susceptibility determinants (**Fig. S4.A**).

### Identification of bacterial genes required for susceptibility to cocktail phages

Following phage challenge, Tn mutants carrying insertions in genes required for successful phage infection were enriched, recovered, and sequenced. Sequencing yielded between 12 and 49 million reads per biological replicate (**Table S3).** For three of the four datasets, 88-95% of reads contained the Himar1 transposon end sequence, and over 99% of these mapped to unique TA sites, resulting in insertion densities of 58-70% and libraries that met the saturation criteria recommended by TRANSIT (i.e., insertion density > 30%, NZ mean > 30, total reads > 1M). One dataset (Phi*Xhv*-2) displayed a lower proportion of Tn-end-containing reads (3%), yielding a reduced insertion density (31%). Although this value was at the lower limit of the recommended range, the insertion density remained sufficient to identify the major susceptibility determinants (**Fig. S3**). Pearson correlation coefficients between biological replicates ranged from 0.67 to 0.94 across all datasets, supporting the reproducibility of the datasets (**Fig. S4.B-D**). These datasets enabled robust statistical comparisons between control and phage-treated conditions (**Table S4**). For each phage, resampling analysis identified genes whose disruption significantly increased bacterial survival during phage challenge as illustrated by the differential fitness volcano plots (log_2_FC > 2, *q-value* < 0.05) (**Fig. S5**). Across the four tested phages, a variable number of enriched genes were found: 9 for Phi*Xhv*-2, 11 for Phi*Xhv*-5, 36 for Phi*Xhv*-7, and 24 for Phi*Xhv*-12, reflecting differences in the complexity of the host determinants required for infection by each phage. These gene lists were then functionally classified using the COG database, revealing three predominant categories: (i) cell wall/membrane/envelope biogenesis, (ii) carbohydrate transport and metabolism, and (iii) intracellular trafficking/secretion (**Fig. S5**). These functional categories are consistent with the central role of bacterial surface structures as primary receptors for phage adsorption and infection. Notably, two major surface structures emerged across multiple datasets: the lipopolysaccharide (LPS) and an outer membrane protein putatively involved in type I secretion (**Fig. 2**). Genes associated with an outer membrane protein and with O-antigen biosynthesis/assembly were significantly enriched for Phi*Xhv*-2 and Phi*Xhv*-5, whereas only LPS-associated gene clusters were significantly enriched for Phi*Xhv*-7 and Phi*Xhv*-12. Two additional significant genes encoded proteins of unknown function (**Table S5**).

**Figure 2.**
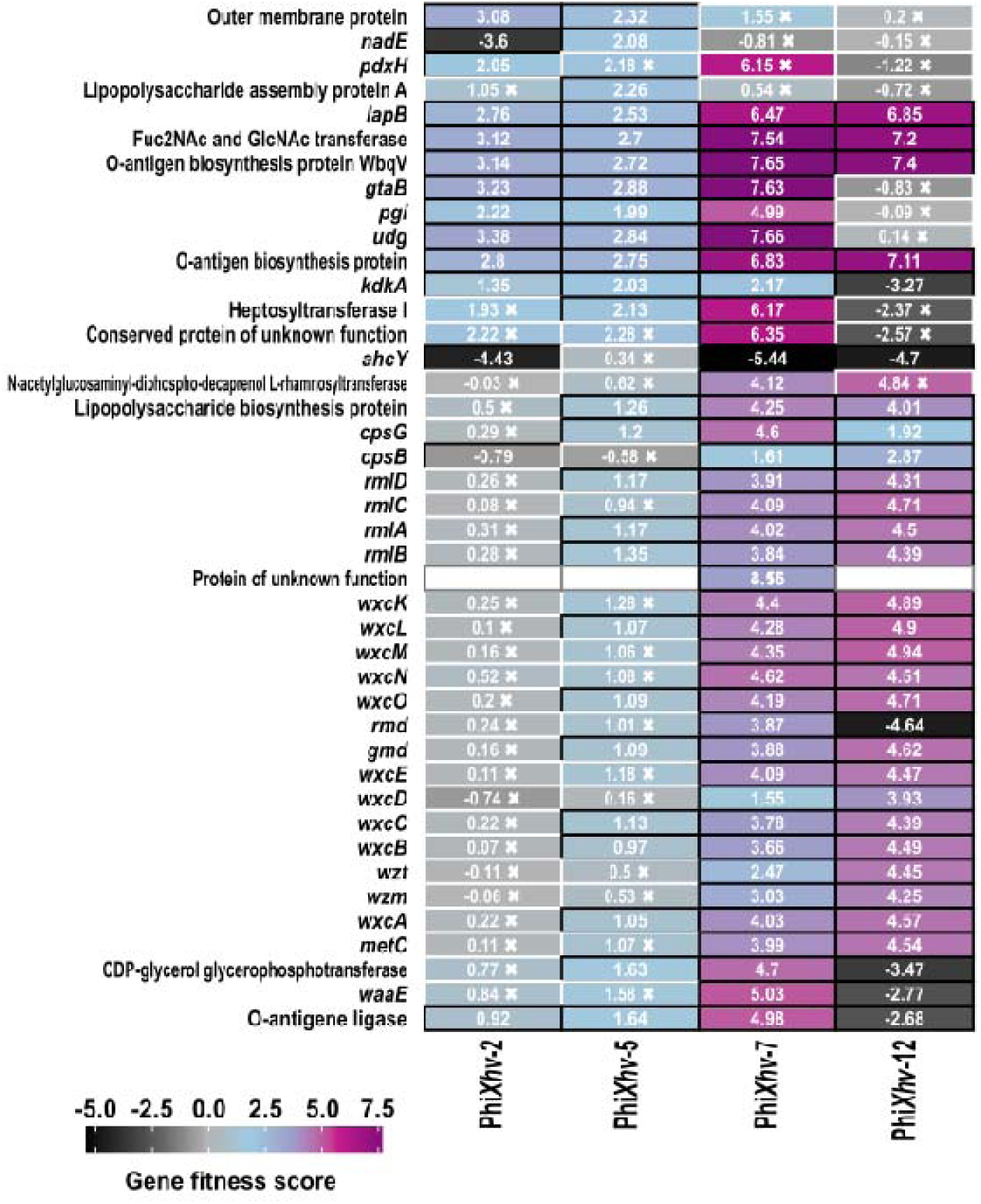
Gene fitness profiles across four bacterium-phage interactions. Heatmap showing gene fitness score (log_2_ fold change) of transposon mutants following selection with phages Phi*Xhv*-2, Phi*Xhv*-5, Phi*Xhv*-7, and Phi*Xhv*-12. Phages Phi*Xhv*-2 and Phi*Xhv*-5 were screened using the CFBP8642 transposon library, Phi*Xhv*-7 using CFBP8641, and Phi*Xhv*-12 using CFBP8638. Genes shown were significantly enriched in at least one phage treatment and are ordered according to their genomic position. Positive fitness scores (fuchsia, purple) indicate enrichment of the corresponding mutants following phage selection, whereas negative scores (black) indicate depletion. Cells outlined in white and marked with a cross indicate non-significant fitness changes (adjusted *p. value* > 0.05).

### Comparative analysis of bacterial determinants for cocktail phage infection and their relevance to plant fitness

The sets of susceptibility determinants identified for Phi*Xhv*-2, Phi*Xhv*-5, Phi*Xhv*-7, and Phi*Xhv*-12 were combined with the previously published dataset for Phi*Xhv*-1 (28) and compared to identify shared and phage-specific determinants of infection (**Fig. 3**). Among the phages infecting ancestral strains (i.e., excluding Phi*Xhv*-28), only four of the 43 identified genes were shared across all phages, all involved in LPS biosynthesis and assembly. In addition, Phi*Xhv*-1, Phi*Xhv*-2, Phi*Xhv*-5, and Phi*Xhv*-7 each required one or two bacterial genes that were unique to its infection profile, potentially involved in receptor modification. Approximately 42% (18/43) of the identified susceptibility determinants were shared exclusively by three phages of the cocktail (Phi*Xhv*-1, Phi*Xhv*-7, and Phi*Xhv*-12), with nearly one-third located within the LPS3 region. Notably, Phi*Xhv*-28, isolated on a mutant defective in the LPS3 region, displayed strict specificity for the altered O-antigen structure and failed to infect ancestral strains. These complementary patterns support distinct infection strategies and reduce the likelihood that a single resistance mutation would compromise cocktail efficacy, thereby justifying the inclusion of six phages in the formulation.

**Figure 3.**
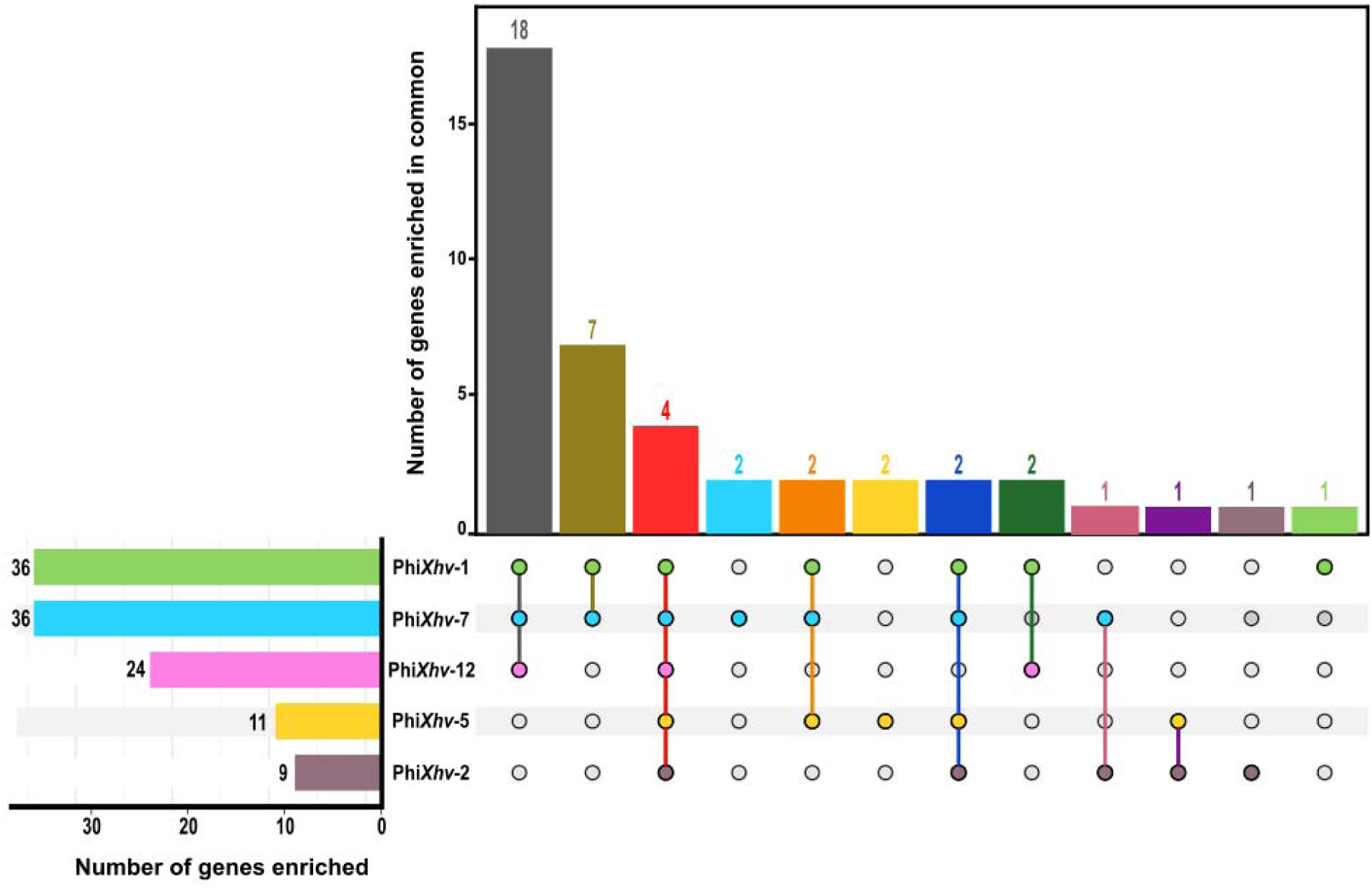
Overlap of *X*. *hortorum* pv. *vitians* genes required for successful infection by five phages of the cocktail. UpSetplot showing the distribution and overlap of significantly enriched genes identified by Tn-seq following selection with Phi*Xhv*-1, Phi*Xhv*-2, Phi*Xhv*-5, Phi*Xhv*-7, and hi*Xhv*-12. Horizontal bars indicate the total number of enriched genes identified for each phage. Vertical bars indicate the number of genes corresponding to each intersection, with colored dots and connecting lines identifying the phage or combination of phages defining each intersection. Genes were considered significantly enriched at log_2_FC > 2 and *q*. *value* < 0.05.

Functional mapping of the identified susceptibility determinants (**Fig. 4**) confirmed the predominance of genes involved in LPS biosynthesis in several phages, together with additional targets related to secretion systems and sugar metabolism pathways. Among the genes identified as critical for phage infection, 63% (27/43) overlapped with genes previously shown to contribute to bacterial fitness *in planta* (34). Strikingly, the four genes required for infection by all tested phages were also key determinants of bacterial colonization and virulence *in planta*. These findings suggest that mutations conferring resistance to multiple phages would simultaneously impair bacterial fitness and virulence *in planta*, thereby increasing the evolutionary robustness of the phage cocktail.

**Figure 4.**
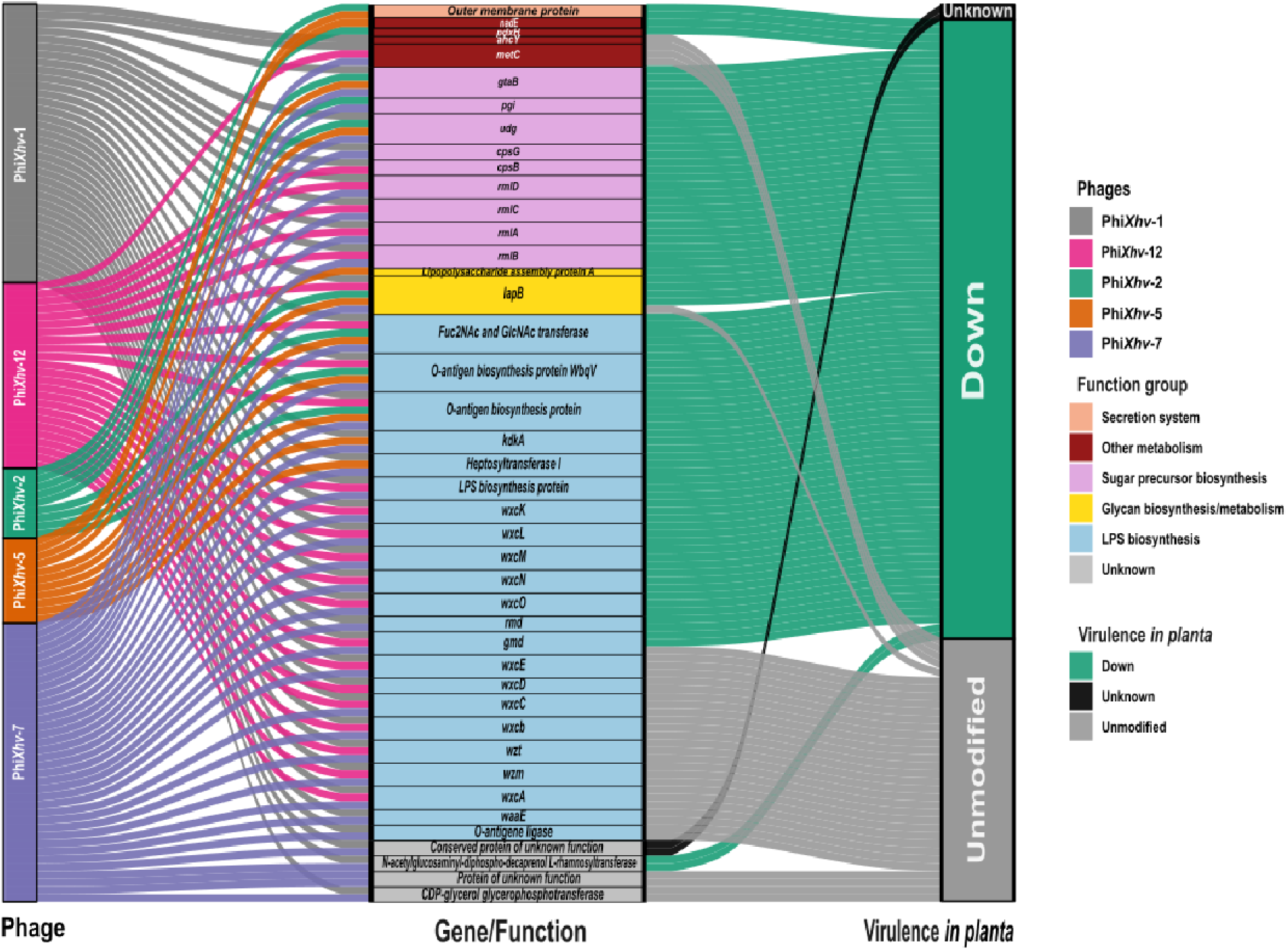
Integration of phage-associated genetic determinants with *in planta* virulence-associated genes. Sankey diagram linking the five phages analyzed by Tn-seq (left) to significantly enriched genes following phage selection (center) and their corresponding classification based on the *in planta* Tn-seq dataset (right) (34). Flow colors the left identify the selecting phage, while gene boxes are color-coded according to predicted functional categories based on KEGG pathways, eggNOG annotations, and KEGG Orthology (KO) identifiers. Functional categories comprise secretion system, other metabolic pathways, sugar precursors biosynthesis, glycan biosynthesis/metabolism, LPS biosynthesis, and genes of unknown functions. Genes were classified according to the *in planta* dataset as associated with reduced fitness/virulence *in planta*, showing no significant change (Unmodified), or not assessed (Unknown).

### Dose and application dependent efficacy of the phage cocktail against bacterial leaf spot of lettuce under growth chamber conditions

Preventive applications of the phage cocktail reduced disease severity in a dose-dependent manner. Disease progression was increasingly delayed with increasing MOI (**Fig. 5.A**), and all phage treatments significantly reduced disease severity compared with the infected control at 20 DPI (**Fig. 5.B).** The extent of disease reduction increased with MOI, whereas the effect of repeated applications depended on the initial MOI, with two applications being more effective than one at MOI 0.1 (*p* = 0.014), but not at MOI 1 (*p* = 0.046).

**Figure 5.**
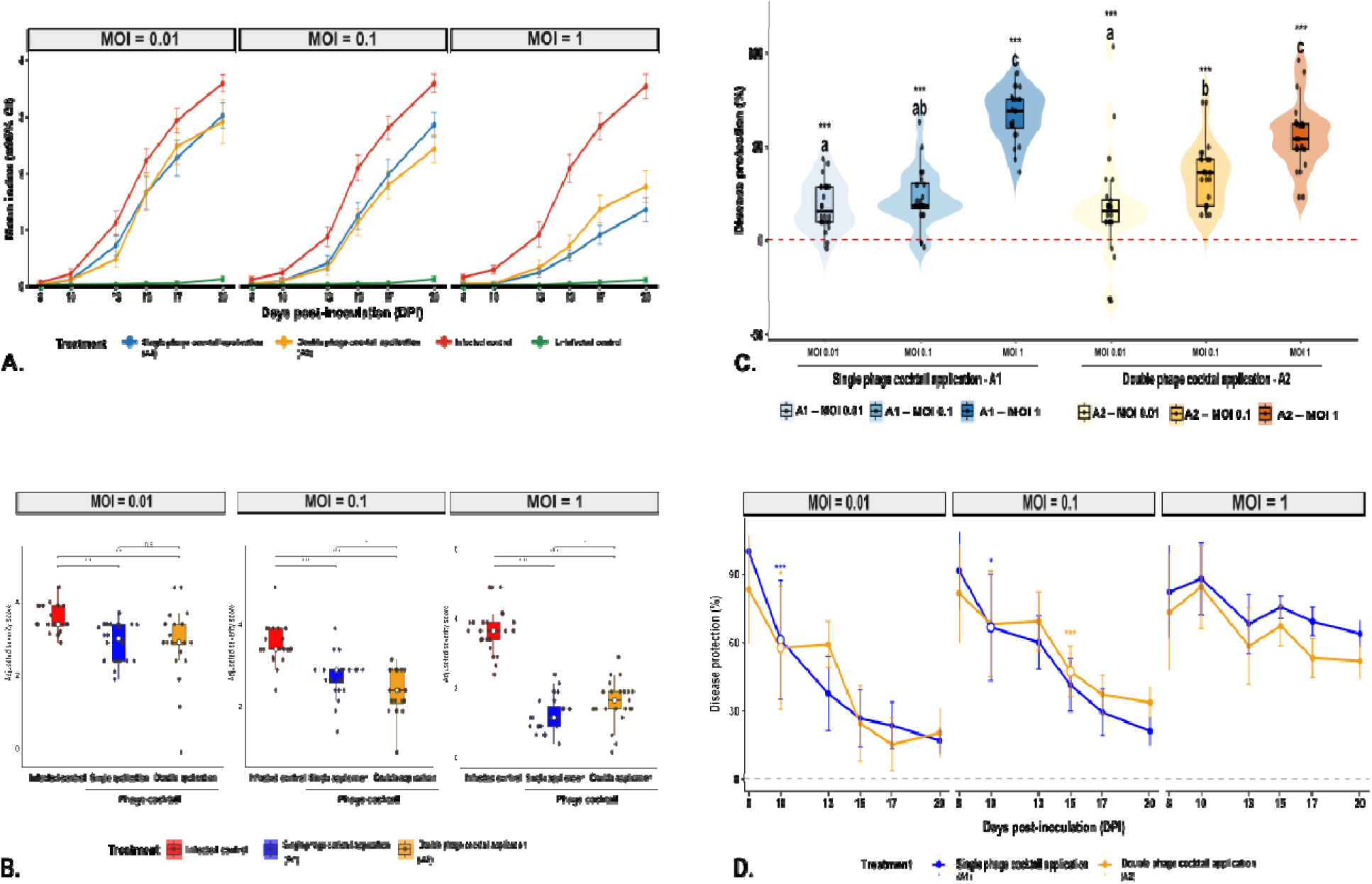
Protective efficacy of the six-phage cocktail against *X*. *hortorum* pv. *vitians* under controlled conditions. **A.** Disease progression from 8 to 20 days post-inoculation (DPI) at multiplicities of infection (MOI) of 0.01, 0.1, and 1. Plants received either a single (A1) or double (A2) phage cocktail application and were compared with infected and uninfected controls. Points represent mean disease severity across three independent experiments and error bar indicate 95% confidence intervals. **B.** Disease severity at 20 DPI for each MOI. Scores were corrected by subtracting the mean of the corresponding uninfected control within each replicate. Violin plots show the distribution of adjusted severity scores per treatment (n=8, 3 independent replicates), embedded boxplots indicate medians and interquartile ranges, and points represent individual plants. Treatments were analyzed separately within each MOI using Gaussian linear mixed-effects models followed by Tukey-adjusted pairwise comparisons. Asterisks indicate significant differences (*p* < 0.05 (*), *p* < 0.01 (**), *p* < 0.001, ns = not significant). **C.** Disease protection (%) conferred by the phage cocktail at 20 DPI according to MOI and number of phage cocktail applications. Protection was calculated from adjusted disease severity relative to the corresponding infected control within each replicate and MOI. Violin plots show individual-plant distributions with embedded boxplots indicating medians and interquartile ranges. Different letters indicate significant differences among phage-treatments combinations across MOI and application schemes (Tukey’s HSD, *p* < 0.05). Asterisks indicate significant differences from the infected control within the corresponding MOI (Dunnett-adjusted contrasts, *p* < 0.001 (***)). **D.** Dynamics of disease protection (%) from 8 to 20 days DPI following single (A1) or double (A2) phage cocktail application at each MOI. Points and error bar represent means and 95% CIs. Within each MOI and application scheme, protection at subsequent assessment dates was compared with the 8-DPI baseline using Sidak-adjusted contrasts. Asterisks indicate significant decreases in protection relative to 8 DPI (*p* < 0.05 (*), *p* <0.01 (**), *p* < 0.001 (***)).

This dose-dependent response was also reflected in disease protection at 20 DPI (**Fig. 5.C**). At MOI 1, both application regimes provided significantly greater protection than all other treatments (*p* < 0.001), with protection reaching 66.2% after one application (A1) and 55.1% after two applications (A2). At MOI 0.1, protection reached 22.2% with A1 and 34.7% with A2, whereas the lowest MOI resulted in 18.1% and 17.2% protection, respectively. Protection remained stable throughout the experiment at MOI 1 (all *p* ≥ 0.087), whereas it declined significantly over time at the lower MOI (**Fig. 5.D**). At MOI 0.1, protection following A2 declined significantly after 14 DPI (Δ = −34.3 ± 9.1, *p* = 8.1 × 10^-4^), while the remaining lower-dose treatments showed a significant decline from 10 DPI onward (all *p* < 0.0297). Overall, these results demonstrate a strong dose-dependent effect of the phage cocktail, whereas the benefit of a second application depended on the initial phage dose.

### Consistent field efficacy of the six-phage cocktail against bacterial leaf spot of lettuce across two growing seasons

Field evaluation of the phage cocktail was conducted during two independent trials performed in summer and autumn 2025. The efficacy of the phage cocktail did not differ between the two trials (likelihood-ratio test for the treatment × season interaction: ^2^ = 0.01, df = 1, *p* = 0.914), indicating a consistent protective effect across growing seasons. In both trials, phage treatment shifted symptom distributions toward lower disease severity classes compared with the infected formulation control (**Fig. 6.A**). Mean disease severity scores (95% confidence intervals) for all treatment conditions are reported in **Table 1**. Compared with the infected formulation control, the phage cocktail reduced the estimated mean symptom severity by 29.9% in summer and 30.6% in autumn (both *p* < 0.0001). These reductions in disease severity translated into improved marketability of harvested lettuce. The probability of producing non-marketable lettuce was significantly reduced by the phage-cocktail (**Fig. 6.B**). Compared with the infected formulation control, the estimated proportion of non-marketable lettuce decreased from 30.7% to 4.7% in summer and from 23.5% to 3.7% in autumn, corresponding to reductions of 84.6% and 84.4%, respectively (both *p* < 0.01). Unexpectedly, the uninfected formulation control exhibited higher symptom severity than the infected formulation control during the autumn trial and also produced greater raw head weights than both formulation-treated conditions. This atypical response may reflect uncontrolled field variability or heterogeneous natural disease pressure under field conditions, although did not affect the estimated efficacy of the phage cocktail relative to the infected formulation control. The effects of the phage cocktail on raw head weight, trimmed head weight, and post-harvest trimming losses were likewise consistent across seasons (likelihood-ratio tests for treatment × season interactions: all *p* ≥ 0.115). Raw head weight remained comparable between the phage treatment and the infected formulation control in both summer (estimated means, 211 and 220 g, respectively; Tukey-adjusted *p* = 0.589) and autumn (360 and 361 g, respectively; *p* = 0.994) (**Fig. 7.A**). Similarly, trimmed head weight did not differ between these treatments in summer (172 and 174 g, respectively; *p* = 0.984) or autumn (310 and 295 g, respectively; *p* = 0.135) (**Fig. 7.B**). Despite the absence of differences in head weight, phage treatment significantly reduced post-harvest trimming losses, which decreased from 20.7% to 18.1% in summer (odds ratio = 0.84, *p* = 0.023) and from 17.8% to 14.0% in autumn (odds ratio = 0.75, *p* = 0.0001) (**Fig. 7.C**). The formulation alone did not exert consistent effects on disease severity or lettuce quality across seasons (**Fig. S6**).

**Figure 6.**
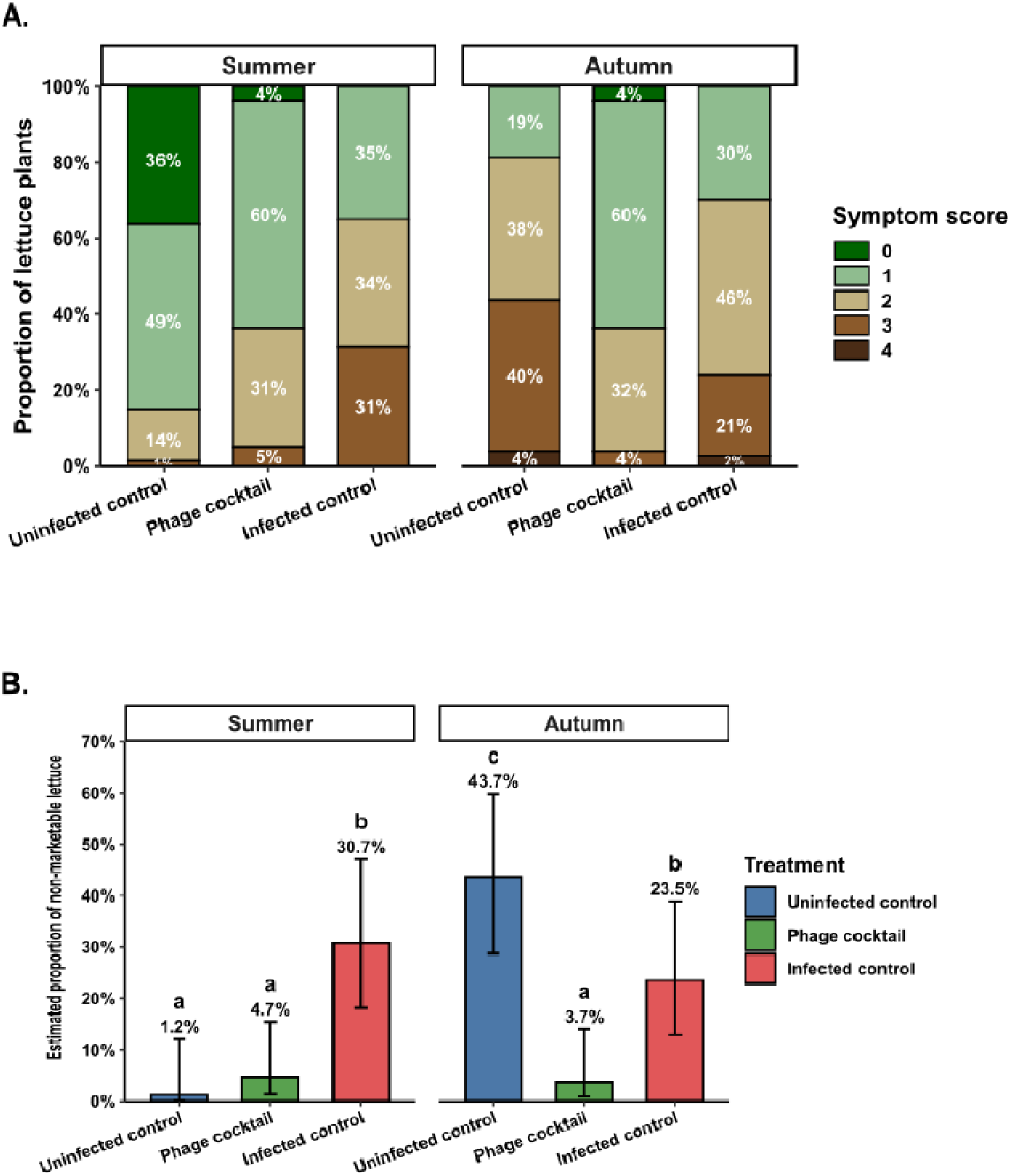
Field efficacy of the six-phage cocktail against bacterial leaf spot of lettuce across two growing seasons. **A.** Distribution of disease severity scores at harvest in the summer and autumn field trials for the uninfected formulation control, phage cocktail treatment, and inoculated formulation control. Stacked bars represent the proportion of lettuce plants in each severity class on a 0-4 scale, with percentages indicated within each segment. Scores ≥ 3 were considered non-marketable. **B.** Model-estimated probability of non-marketable lettuce for each treatment during the summer and autumn field trials. Bars represent estimated probabilities and errors bars indicate 95% confidence intervals. Data were analyzed using a binomial generalized linear mixed-effects model including treatment, seasons, and their interaction as fixed effects and field plot nested within season as a random effect. Different letters indicate significant differences among treatments within each season based on Tukey-adjusted pairwise comparison (*p* <0.05).

**Figure 7.**
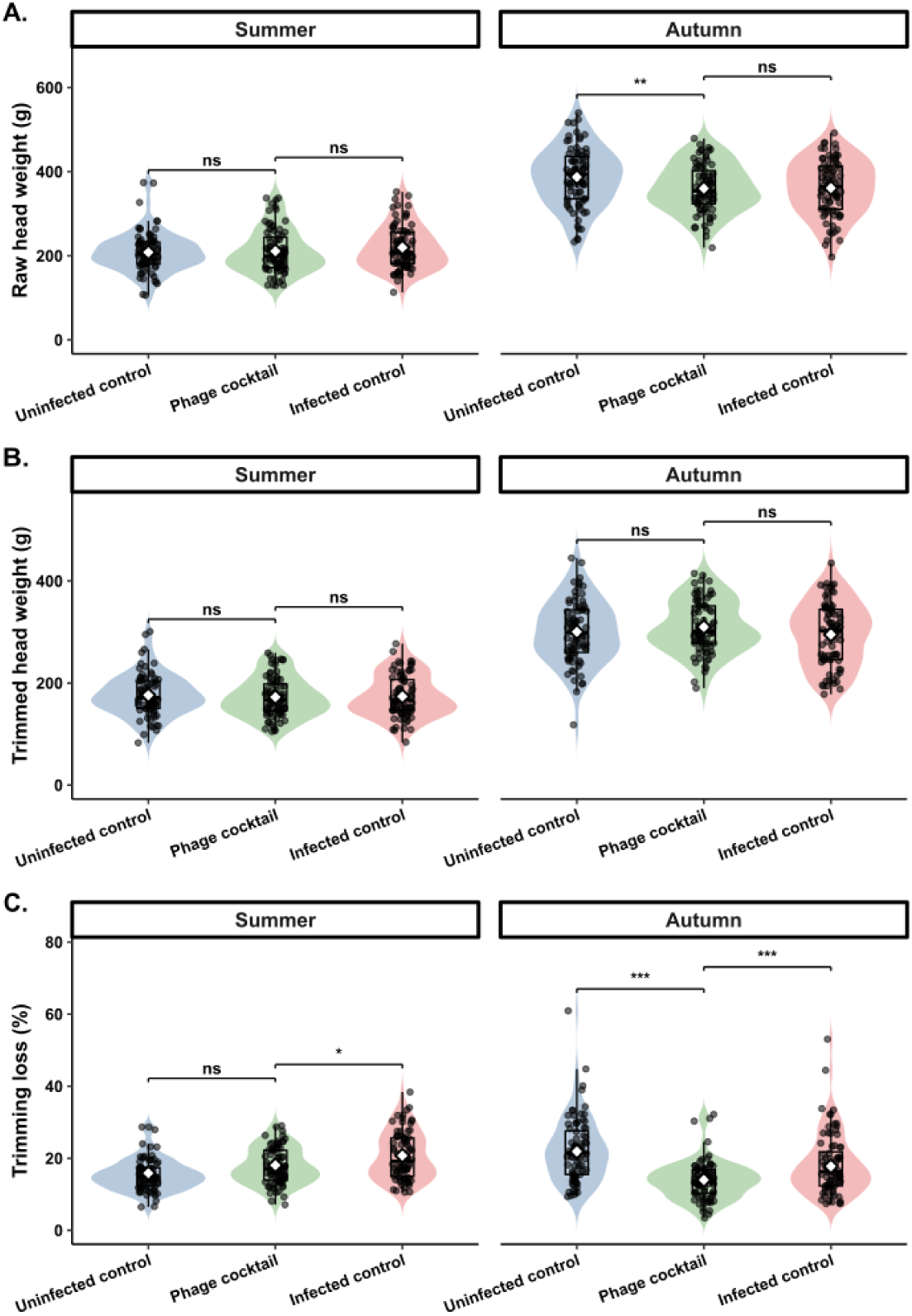
Effects of the six-phage cocktail on lettuce yield and trimming losses across two growing seasons. **A.** Raw head weight at harvest. **B.** Trimmed head weight after removal of damaged outer leaves. **C.** Trimming loss, expressed as a percentage of raw head weight. Measurements are shown for the uninoculated formulation control, phage cocktail treatment, and inoculated formulation control in the summer and autumn field trials. Violin plots represent the distribution of individual plant values (dots), with embedded boxplots indicating medians and interquartile ranges. Pairwise differences between treatments within each season are indicated by asterisks (*p* < 0.05 (*), *p* <0.01 (**), *p* < 0.001 (***), *p* ≥ 0.05 (ns)) following Tukey-adjusted pairwise comparisons.

**Table 1.** Mean symptom severity scores across the two field trials.

| Season | Treatment | Observed mean symptom score (95% IC) |
| --- | --- | --- |
| Summer | Uninfected control | 0.61 (0.10-1.12) |
|  | Uninfected formulation control | 0.80 (0.65-0.95) |
|  | Phage cocktail | 1.38 (0.70-2.05) |
|  | Infected control | 1.50 (0.89-2.11) |
|  | Infected formulation control | 1.96 (1.23-2.70) |
| Autumn | Uninfected control | 2.60 (2.03-3.17) |
|  | Uninfected formulation control | 2.29 (1.74-2.83) |
|  | Phage cocktail | 1.36 (0.69-2.03) |
|  | Infected control | 1.94 (1.67-2.20) |
|  | Infected formulation control | 1.96 (1.27-2.66) |

Values are means of plot-level mean symptom scores (n = 4 plots per treatment and season; 20 plants per plot). Disease severity was scored on a 0–4 ordinal scale; 95% confidence intervals were calculated from plot-level means. Statistical inference was based on the ordinal mixed-effects model described in Materials and Methods.

## DISCUSSION

Phage biocontrol is considered one of the most promising strategies for the biological control of bacterial plant diseases. However, its successful implementation depends on the rational design of phage cocktails combining complementary host ranges and distinct infection strategies to ensure efficacy, durability, and field applicability. In this study, we formulated and evaluated a six-phage cocktail targeting *Xanthomonas hortorum* pv. *vitians*, the causative agent of bacterial leaf spot of lettuce.

While the selection of phages with characteristics adapted to biocontrol is essential (35), no universally accepted guidelines exist for the rational assembly of phage cocktails targeting bacterial pathogens. A key consideration lies in the interplay between phage-bacteria and phage-phage interactions, which can significantly influence the overall efficacy of a phage cocktail. Although combining phages is generally intended to broaden the host range and mitigate resistance emergence, some studies have reported that phage mixtures can exhibit interference effects, where the lytic activity of the cocktail is reduced compared to the theoretical sum of its individual components (36). For instance, Bourdin *et al*. (37) found that a combination of three phage T4-like phages resulted in a narrower host range against *E*. *coli* strains than when the phages were applied individually. The precise mechanisms underlying antagonist interactions are still poorly characterized, but competition during co-infection, superinfection exclusion, or shared receptor usage are frequently cited as potential causes (38, 39). These findings underscore the importance of selecting phages that rely on distinct infection mechanisms or bacterial determinants, thereby reducing the risk of mutual interference and improving the breadth of bacterial targeting. In our study, this strategy proved effective. Host range analysis of the six-phage cocktail revealed no detectable interference among phages: strains lysed by a single phage remained susceptible within the cocktail, and the overall lytic spectrum matched the theoretical sum of the individual profiles. This preservation of individual activity likely reflects the diversity of receptor usage among the selected phages.

The diversity of bacterial determinants involved in phage infection revealed three distinct infection strategies within the phage cocktail. First, approximately 50% and 75% of the total genes identified for Phi*Xhv*-7 and Phi*Xhv*-12, respectively, were common to Phi*Xhv*-1 and include key components of the LPS biosynthesis pathway. These findings support the hypothesis that these three phages target LPS structures, consistent with previous findings on Phi*Xhv*-1 (28). Second, Phi*Xhv*-28 displays a highly restricted host range and only infects strains with a modified O-antigen structure as described previously (29). Finally, phages Phi*Xhv*-2 and Phi*Xhv*-5 likely rely on dual recognition of the O-antigen and an outer membrane protein, predicted to belong to the type I secretion system. To our knowledge, this is the first report suggesting a secretion system component might serve as a phage receptor in *Xanthomonas*. Similar strategies have been identified in other bacterial systems, where components of the type II secretion system (T2SS) have been identified as phage receptors in *Ralstonia solanacearum* (40) and *Vibrio cholerae* (41). Moreover, sequential recognition of the O-antigen followed by a secondary interaction with an outer membrane protein, OmpA or OmpC, has been reported for phage Sf6 infecting *Shigella flexneri* (42, 43). Consistent with this hypothesis, several studies have shown that phage cocktails using different receptors (between 2 and 3) delayed the emergence of resistant mutants and did not develop cross-resistance (44, 45).

Beyond the role of phage receptors, components of the type 1 secretion system (T1SS) in *Xanthomonas* are known to contribute to the early stages of plant colonization. Specifically, they are involved in the secretion of bacteriocin, proteins with non-peptide RTX motifs and large adhesin, all of which promote bacterial attachment to host surfaces (46). Mutation of the outer membrane protein identified as a putative receptor in our study has previously been associated with reduced *in planta* fitness (34). More broadly, 63% of the genes required for phage infection in the cocktail were also found to contribute to bacterial survival *in planta*. Such fitness costs have already been demonstrated for the receptors of Phi*Xhv*-1, which were associated with reduced motility and virulence (28). These observations highlight the evolutionary trade-offs associated with resistance. A study by Wang *et al*. (2019) demonstrated that increasing the number of phages in a cocktail progressively reduced the fitness of *Ralstonia solanacearum* mutants resistant to phage infection. The development of resistance to a three-phage cocktail resulted in a marked reduction in bacterial growth and competitiveness, compared to resistance against one or two phages, suggesting that multi-phage resistance enhances the associated fitness costs and evolutionary trade-offs. In another study, the combined effect of trade-offs and repeated phage applications was so strong that no resistant clones could be recovered after treatment with a phage cocktail (47). These findings are consistent with the expected robustness of well-designed cocktails when phages exploit functionally diverse receptors that are also involved in other key bacterial traits. Despite its efficacy, phage cocktail biocontrol in the phyllosphere must overcome numerous environmental and biological constraints that influence treatment success. Indeed, phage-mediated protection against bacterial plant diseases has been shown to depend on the timing, frequency, and density of application. Under controlled growth-chamber conditions, the highest phage concentration (10^8^ PFU.mL^-1^) applied prior to pathogen inoculation provided the most effective suppression of disease symptoms. However, even the lowest dose tested (10^6^ PFU.mL^-1^) still led to a significant reduction in symptom severity under controlled conditions, indicating that protective effects can be achieved across a broad range of phage concentrations. These observations are consistent with previous studies reporting effective disease suppression with phage concentrations ranging from 10^5^ to 10^8^ PFU.mL^-1^ (26), although the dose-response relationship is not always strictly linear (48). For example, Balogh *et al*. (48) observed similar levels of protection against *Xanthomonas campestris* pv. *vesicatoria* using 10^6^ and 10^8^ PFU.mL^-1^, whereas 10^4^ PFU.mL^-1^ was ineffective. Protection is generally attributed to the reduction of pathogen populations following phage replication. For instance, the preventive foliar application of the phage Medea1 against *Pseudomonas syringae* pv. *tomato* resulted in a ∼60-fold reduction in bacterial density on leaves within 24 hours (49). Likewise, Wang *et al*. (47) found that repeated phage treatments progressively reduced *Ralstonia solanacearum* populations in the rhizosphere, with the most consistent suppression observed after three applications. This improved efficacy coincided with the maintenance of higher phage titers, whereas phage populations declined rapidly following one or two applications. In addition to direct bacterial lysis, some studies suggest that phages may also reduce disease symptoms through indirect mechanisms. Notably, Skliros *et al*., (49) reported that phage application enhanced the expression of salicylic acid-dependent defense genes in tomato, suggesting that phages might also stimulate plant immune responses under certain conditions. When applied to the phyllosphere, however, phages are exposed to multiple environmental and biological constraints. Among them, UV radiation (50) and low surface moisture (51) have been shown to reduce phage persistence, limit their diffusion and interaction with bacterial hosts. To address these restrictions, we included both clay (bentonite) and a sticker adjuvant in the foliar formulation, which may help increase phage persistence on the leaf surface. Supporting this hypothesis, *in vitro* studies using kaolin, a structural analogue of bentonite, have demonstrated improved phage survival under abiotic stress. Additionally, Magar *et al* (52) reported that the addition of an adjuvant in their phage formulation improved disease control, suggesting a potential benefit of such additives. Although the formulation alone did not consistently reduce disease severity across the two field trials, it likely contributed to phage persistence on the leaf surface, which may have enhanced the efficacy of the phage cocktail. Another important limitation of foliar applications is the apparent lack of systemic phage movement within plant tissues. Unlike root-applied phages, which can be translocated into stems and leaves (53, 54), foliar-applied phages have never been recovered from internal plant tissues. This lack of systemic movement likely limits their capacity to reach bacterial population that have already colonized internal plant compartments, a critical consideration for *Xanthomonas* spp. which are known to rapidly transition from an epiphytic to an endophytic lifestyle in infection condition (55). In our study, the limited and transient protection observed at lower phage doses (4 × 10^6^ PFU.mL^-1^) or after single application of an intermediate phage concentration (4 × 10^7^ PFU.mL^-1^), may be explained by these limitations. Across two independent field trials conducted during contrasting growing seasons, the phage cocktail consistently reduced symptom severity by approximately 30% following three weekly applications of the phage cocktail. This level of protection is comparable those reported against other *Xanthomonas* diseases under field conditions, including *X*. *campestris* pv. *campestris* (56) and *X. axonopodis* pv. *alii* (57). In the later study, phages remained detectable on the leaf surface for up to four days post-application, suggesting that the interval between treatments should ideally be adjusted to match this persistence window in order to enhance protection. Several strategies have been proposed to enhance the efficacy of phage-based biocontrol. Integrated pest management combining phages with competitive antagonistic bacteria have shown promising results. For instance, in the context of *Ralstonia solanacearum*, the co-application of a lytic phage and a *Bacillus amyloliquefaciens* strain resulted in a synergistic effect, achieving up to 40% disease protection. This effect was attributed to a trade-off whereby phage-resistant bacterial mutants became more susceptible to antibiotic compounds produced by the bacteria (58). Similarly, co-treatment with *Pseudomonas lalkuanensis*, a plant growth-promoting rhizobacterium, and phages improved seedling emergence, enhanced plant growth, and provided better disease suppression, while also reducing soil salinity (59). In addition, the use of plant defense activators such as acibenzolar-*S*-methyl in combination with phages led to further reductions in disease severity. However, in this case, no significant improvement was observed in the yield of marketable fruit (60). By contrast, across two independent field trials, phage treatment not only consistently reduced disease severity but also improved lettuce marketability. Although raw and trimmed head weights were not consistently increased, phage-treated lettuce exhibited significantly lower trimming losses and a marked reduction in the proportion of nonmarketable heads, demonstrating that disease suppression translated into tangible post-harvest benefits. Similar agronomic benefits of phage biocontrol have also been observed in other systems. For example, in field trials targeting *Dickeya solani*, phage application led to a 13% increase in tuber yield. This improvement was largely attributed to an increase in tuber size rather than number (61). In a related study, yield gains ranged from 4 to 31.9% depending on infection severity, with the highest increases observed under mild infection conditions when phage treatment was applied (62). Together with our results, these studies highlight the potential of phage biocontrol to improve not only disease management but also the agronomic value of harvested crops.

## CONCLUSION

This study demonstrates the potential of a rationally designed multireceptor phage cocktail for the control of bacterial leaf spot of lettuce caused by *Xanthomonas hortorum* pv. *vitians*. By combining six well-characterized phages with complementary host ranges and distinct infection strategies, we achieved broad coverage of *X. hortorum* pv. *vitians* while maintaining strict pathovar specificity. The identification of distinct bacterial determinants involved in phage infection, including structures associated with virulence and *in planta* colonization, provided a mechanistic rationale design of the phage cocktail. Under controlled growth-chamber conditions, this rationally designed cocktail provided dose-dependent protection against bacterial leaf spot, and was subsequently validated under field conditions. Across two independent field trials in contrasting growing seasons, phage treatment consistently reduced disease severity and substantially improved lettuce marketability while reducing post-harvest trimming losses. The absence of consistent effects from the formulation alone further supports the contribution of phage activity to the observed disease control. These findings provide the first field demonstration of an effective biocontrol solution against *X*. *hortorum* pv. *vitians* and highlight the value of integrating complementary host ranges and infection mechanisms into the rational design of phage cocktails. Future studies should whether this multireceptor strategy enhances the evolutionary durability of phage cocktails by monitoring resistance emergence and phage-pathogen population dynamics *in planta* over successive cropping cycles.

## Acknowledgments

The authors are grateful to Julie Baltenneck and Erwan Gueguen for their help and expertise with the Tn-seq experiments. We also thank Nicolas Taveau for his contribution to the preparation and large-scale production of the phage cocktail used in the field trials. We are especially grateful to Florian Calabro for his valuable support during the establishment and monitoring of the field trials. We also express our gratitude to the FNX and Phages.fr networks for their insightful discussions during scientific meetings. We thank the greenhouse and growth-chamber platform of the FR BioSciences (Université Claude Bernard Lyon 1) for providing access to their facilities and technical support, particularly Elise Lacroix and Lilou Piegay for their assistance during the experiments. Additionally, we acknowledge the sequencing and bioinformatics expertise of the I2BC High-Throughput Sequencing Facility, supported by France Génomique (funded by the French National Program “Investissement d’Avenir” ANR-10-INBS-09).

## Author contributions

A.BA. designed the study, conducted the experiments, curated and analyzed the data, generated the visualizations, and wrote the original draft. I.R. participated in all experimental work and contributed to data analysis. D.A. assisted with Tn-seq data analysis. H.A. developed and evaluated preliminary phage cocktail prototypes that guided the selection of the final cocktail. C.A. contributed to the construction of the Tn-seq libraries and assisted with the phage challenge experiments. D.C. contributed to methodology development, provided scientific guidance on phage cocktail formulation, and supplied the phage stocks for field applications. M.G.B. coordinated R&D activities at GREENPHAGE and contributed to experimental design. A.BU. coordinated the field trial, including planting, phage applications, and disease assessments. F.B. conceptualized and supervised the study, secured funding and managed the overall project. All authors reviewed and edited the manuscript.

## Data availability statement

Transposon insertion sequencing raw reads have been deposited in the NCBI SRA database under accession numbers SRX29880918 to SRX29880931 and are associated with BioProject no. PRJNA1297948. Data supporting the conclusions of this article are available within the article and its Supplementary Information files. Additional experimental datasets have been deposited in Zenodo (DOI: 10.5281/zenodo.21991557, (1)) and are available to editors and reviewers through a private access link: https://zenodo.org/records/21991557?preview=1&token=eyJhbGciOiJIUzUxMiJ9.eyJpZCI6ImUxMGY2NWZjLTBlOWEtNDMzYy05NTgxLWMzYmI3MjY4NzVkNCIsImRhdGEiOnt9LCJyYW5kb20iOiI0OGE2NmM5OTY2ODNkNDdiZTU4ZDFjNzFhZWQ2ODQzNiJ9.OQODXIxNK8NW5sXQloJIn6cn_GOK5q1aRhJV8h26efr-glYcqE0rfsoXqq4c6OVtbqlrtNObXzcJ54O9a3oIbA. The Zenodo repository will be made publicly available upon publication.

## Supplementary tables

**Table S1.** List of bacterial strains used in this study.

**Table S2.** Long-term stability of six individual phages and the corresponding six-phage cocktail after 12 months of storage at 4°C.

**Table S3.** Transposon-insertion sequencing (Tn-seq) statistics before Totreads normalization.

**Table S4.** Unfiltered Tn-seq dataset: differential analysis between phage and control conditions following resampling (1) Phi*Xhv*-2 vs. control library CFBP8642, (2) Phi*Xhv*-5 vs. control library CFBP8642, (3) Phi*Xhv*-7 vs. control library CFBP8641, and (4) Phi*Xhv*-12 vs. control library CFBP8638.

**Table S5.** Genes required for infection by four phages identified by Tn-seq.

## Supplementary figures

**Figure S1. Qualitative disease severity scale for bacterial leaf spot of lettuce, adapted from Morinière *et al*.** (33). Disease severity scores range from 0 to 5: 0= no visible symptom, 1= fewer than 10 isolated necrotic spots (<2 mm), 2= more than 10 isolated necrotic spots (<2 mm), 3= coalescing lesions (<1 cm) on at least two leaves, 4= large coalescing lesions (>1 cm), 5= extensive necrosis covering at least one third of the surface on two or more leaves. Representative photographs illustrate each severity class.

**Figure S2.** Environmental conditions, disease severity scoring scale, and experimental design of the field trials evaluating the six-phage cocktail against *X*. *hortorum* pv. *vitians*. A. Weather conditions during the summer and autumn field trial at the CTIFL experimental station in Brindas, France. Daily precipitation and irrigation are shown as bars. Lines represent daily mean temperature and relative humidity, and vertical lines indicate the corresponding daily minimum-to-maximum ranges. **B.** Disease severity scoring scale used to assess bacterial leaf spot symptoms on lettuce at harvest. Representative photographs illustrate the five severity classes (CL0-CL4), ranging from asymptomatic plants (CL0) to plants with severe disease symptom (CL4). Plants assigned to CL0-CL2 were considered marketable, whereas plants assigned to CL3-CL4 were considered non-marketable. **C.** Field layout showing the randomized complete block design and spatial distribution of the experimental treatments. Treatments included the six-phage formulated cocktail, infected and uninfected controls, and their corresponding formulation controls. Each treatment was replicated across four field plots of 7.5 m^2^ (80 plants per plot), totaling 320 plants per treatment. Plot dimensions, orientation and areas of contaminated and uncontaminated soil are indicated.

**Figure S3. Genome-wide transposon insertion profiles on Tn-seq libraries before and after phage challenge.** Transposon insertion counts are plotted according to their genomic coordinates for each *X*. *hortorum* pv. *vitians* strain. **A.** Genome-wide insertion density across the genome for the novel transposon mutant libraries of strains CFBP8641 and CFBP8642. (**B-D)** Genome-wide insertion profiles of untreated controls and corresponding phage-treated libraries. **B.** CFBP8638 control and CFBP8638 library following challenge with Phi*Xhv*-12. **C.** CFBP8641 control and CFBP8641 library following challenge with Phi*Xhv*-7. **D.** CFBP8642 control and CFBP8642 library following challenge with Phi*Xhv*-2 or Phi*Xhv*-5. Untreated controls were grown in TSB without phage.

**Figure S4. Biological reproducibility of Tn-seq datasets.** Pairwise correlations of gene-level read counts between two biological replicates are shown for each experimental condition. Each dot represents a gene, with axes indicating the read count obtained for that gene in biological replicate 1 and biological replicate 2. Pearson correlation coefficients (*r*) and associated *p. values* are indicated in each plot. The red dashed line indicates the identity line (y=x). **A.** Correlation plots for the novel transposon mutant libraries of strains CFBP8641 and CFBP8642. (**B-D)** Correlation plots for untreated controls and corresponding phage-treated libraries. **B.** CFBP8638 control and CFBP8638 library following challenge with Phi*Xhv*-12. **C.** CFBP8641 control and CFBP8641 library following challenge with Phi*Xhv*-7. **D.** CFBP8642 control and CFBP8642 library following challenge with Phi*Xhv*-2 or Phi*Xhv*-5. Untreated controls were grown in TSB without phage.

**Figure S5.** Identification and functional classification of *X*. *hortorum* pv. *vitians* genes associated with successful infection by four phages of the cocktail using Tn-seq. Volcano plots show the differential abundance of transposon mutants between untreated control and phage-treated libraries. Significantly enriched genes (log_2_FC > 2, *q*.*value* < 0.05) are highlighted in orange, and the number of significantly enriched genes (*n*) is indicated for each dataset. Dashed lines indicate the significance and log_2_FC thresholds used to identify enriched genes. Corresponding bar plots show the distribution of significantly enriched genes among functional categories based on Clusters of Orthologous Groups (COG) annotations. (**A**) Phi*Xhv*-2, screened against the CFBP8642 transposon library, (**B**) Phi*Xhv*-5, screened against the CFBP8642 transposon library, (**C**) Phi*Xhv*-7, screened against the CFBP8641 transposon library, (**D**) Phi*Xhv*-12, screened against the CFBP8638 transposon library.

**Figure S6. Effect of the formulation on disease severity and yield-related traits in the field trials. A.** Distribution of disease severity scores at harvest in uninoculated and inoculated controls, with or without formulation, during the summer and autumn field trials. Stacked bars represent the proportion of lettuce plants in each disease severity class (0-4), with percentages indicated within each segment. **B.** Raw head weight at harvest. **C.** Trimmed head weight after removal of damaged outer leaves. **D.** Trimming loss, express as a percentage of raw head weight. For each parameter, formulation and no-formulation treatments are compared separately for uninoculated and inoculated controls in each season. Violin plots show the distribution of individual plant values (dots), with embedded boxplots indicating medians and interquartile ranges. Asterisks indicate significant differences between formulation and no-formulation treatments (*p* < 0.05 (*), *p* < 0.01 (**), *p* < 0.001, ns = not significant).

