## Supplementary figures for "A multireceptor six-phage cocktail consistently controls bacterial leaf spot of lettuce caused by *Xanthomonas hortorum* pv. *vitians* and improves harvest quality"

**Running title: Phage cocktail biocontrol of *X*. *hortorum* pv. *vitians*.**

Anaelle Baud^1,#^, Inès Rougis^1^, Danis Abrouk^1^, Hanane Amari^3^, Clara Aubremaire^1^, Denis Costechareyre^3^, Marie Graindorge Beaume^3^, Alexandre Burlet^2^, Franck Bertolla^1,#^

**^1^ Université Lyon 1**, CNRS, INRAE, LEM, UMR 5557, UMR 1418 Villeurbanne, France.

**^2^ Centre Technique Interprofessionnel des Fruits et Légumes (CTIFL)**, Antenne de Brindas, 69126 Brindas, France

**^3^ GREENPHAGE**, 34830 Clapiers, France.

**Content:**

**Supplementary figures**

**Figure S1.** Qualitative disease severity scale for bacterial leaf spot of lettuce, adapted from Morinière *et al*.

**Figure S2.** Environmental conditions, disease severity scoring scale, and experimental design of the field trials evaluating the six-phage cocktail against *X*. *hortorum* pv. *vitians*.

**Figure S3.** Genome-wide transposon insertion profiles on Tn-seq libraries before and after phage challenge.

**Figure S4.** Biological reproducibility of Tn-seq datasets.

**Figure S5.** Identification and functional classification of *X*. *hortorum* pv. *vitians* genes associated with successful infection by four phages of the cocktail using Tn-seq.

**Figure S6.** Effect of the formulation on disease severity and yield-related traits in the field trials.

**Reference**

**Supplementary figures**

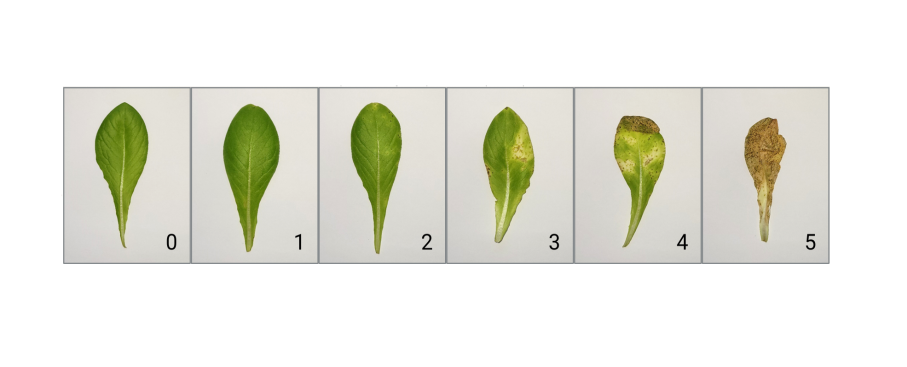

**Figure S1. Qualitative disease severity scale for bacterial leaf spot of lettuce, adapted from Morinière *et al*.** (1)**.** Disease severity scores range from 0 to 5: 0= no visible symptom, 1= fewer than 10 isolated necrotic spots (<2 mm), 2= more than 10 isolated necrotic spots (<2 mm), 3= coalescing lesions (<1 cm) on at least two leaves, 4= large coalescing lesions (>1 cm), 5= extensive necrosis covering at least one third of the surface on two or more leaves. Representative photographs illustrate each severity class.

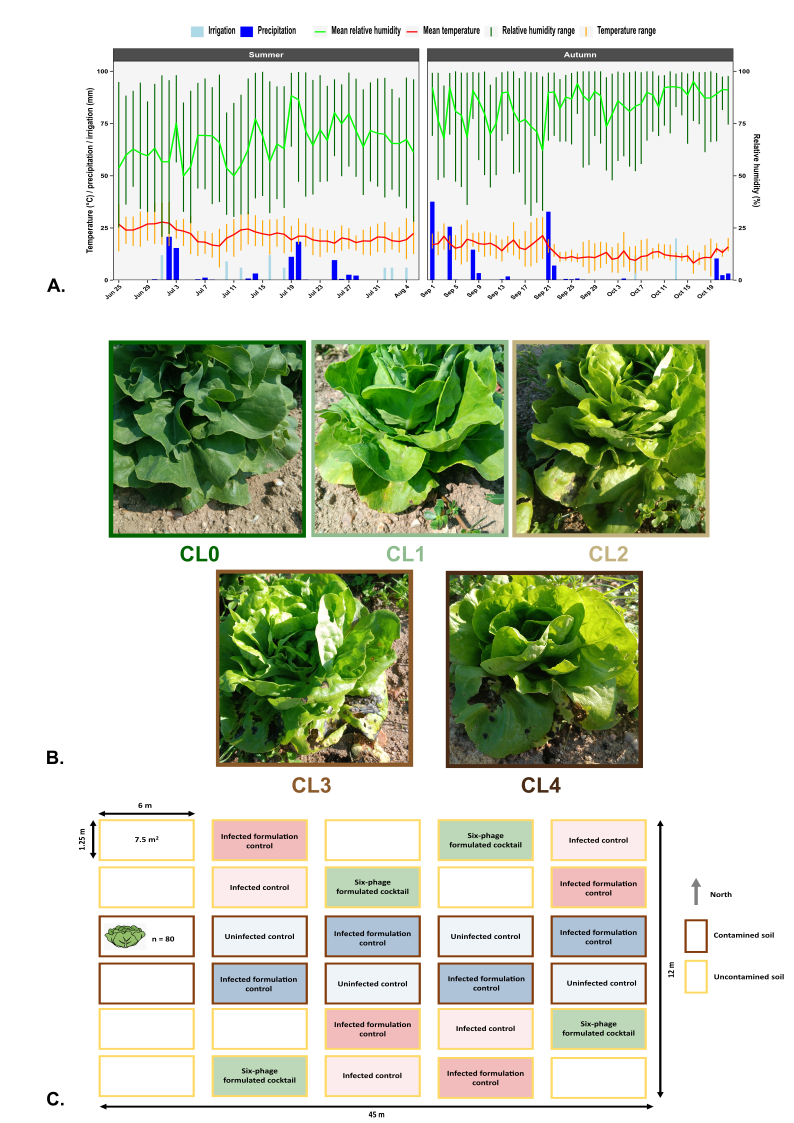

**Figure S2. Environmental conditions, disease severity scoring scale, and experimental design of the field trials evaluating the six-phage cocktail against *X*. *hortorum* pv. *vitians*. A.** Weather conditions during the summer and autumn field trial at the CTIFL experimental station in Brindas, France. Daily precipitation and irrigation are shown as bars. Lines represent daily mean temperature and relative humidity, and vertical lines indicate the corresponding daily minimum-to-maximum ranges. **B.** Disease severity scoring scale used to assess bacterial leaf spot symptoms on lettuce at harvest. Representative photographs illustrate the five severity classes (CL0-CL4), ranging from asymptomatic plants (CL0) to plants with severe disease symptom (CL4). Plants assigned to CL0-CL2 were considered marketable, whereas plants assigned to CL3-CL4 were considered non-marketable. **C.** Field layout showing the randomized complete block design and spatial distribution of the experimental treatments. Treatments included the six-phage formulated cocktail, infected and uninfected controls, and their corresponding formulation controls. Each treatment was replicated across four field plots of 7.5 m2 (80 plants per plot), totaling 320 plants per treatment. Plot dimensions, orientation and areas of contaminated and uncontaminated soil are indicated.

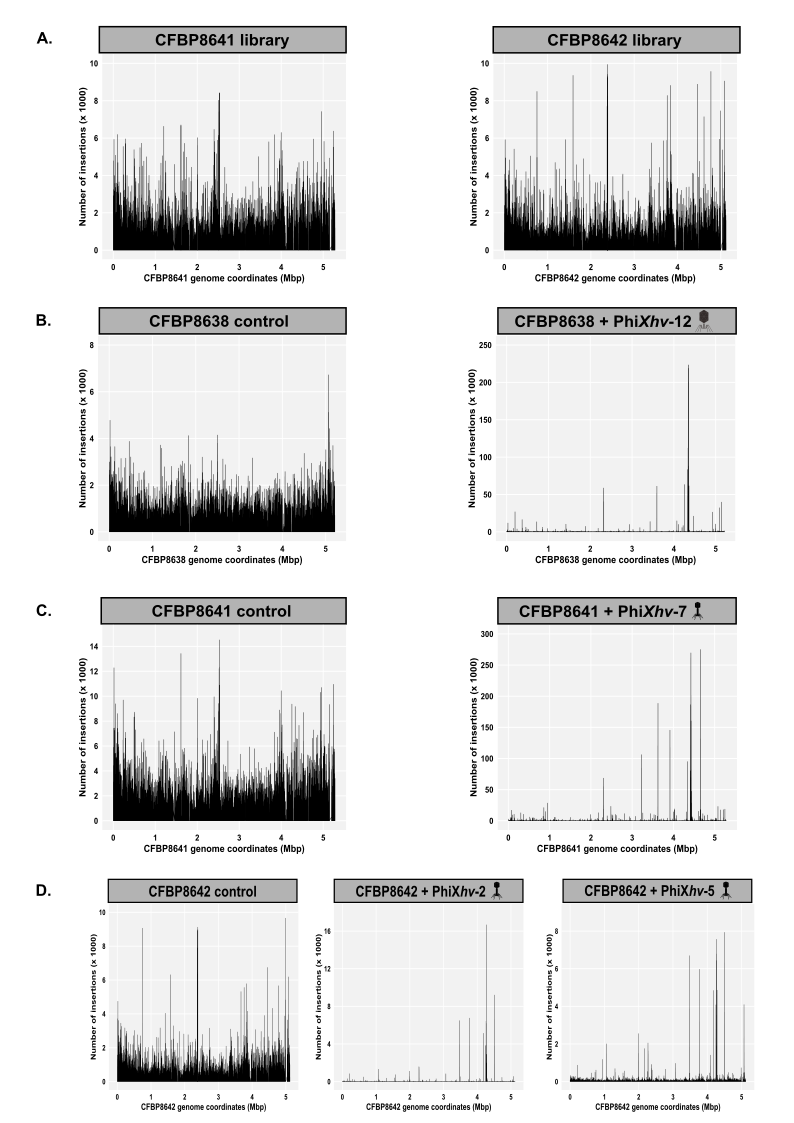

**Figure S3. Genome-wide transposon insertion profiles on Tn-seq libraries before and after phage challenge. Transposon insertion counts are plotted according to their genomic coordinates for each *X*. *hortorum* pv*. vitians* strain.** **A.** Genome-wide insertion density across the genome for the novel transposon mutant libraries of strains CFBP8641 and CFBP8642. **(B-D)** Genome-wide insertion profiles of untreated controls and corresponding phage-treated libraries. **B**. CFBP8638 control and CFBP8638 library following challenge with Phi*Xhv*-12. **C.** CFBP8641 control and CFBP8641 library following challenge with Phi*Xhv*-7. **D.** CFBP8642 control and CFBP8642 library following challenge with Phi*Xhv*-2 or Phi*Xhv*-5. Untreated controls were grown in TSB without phage.

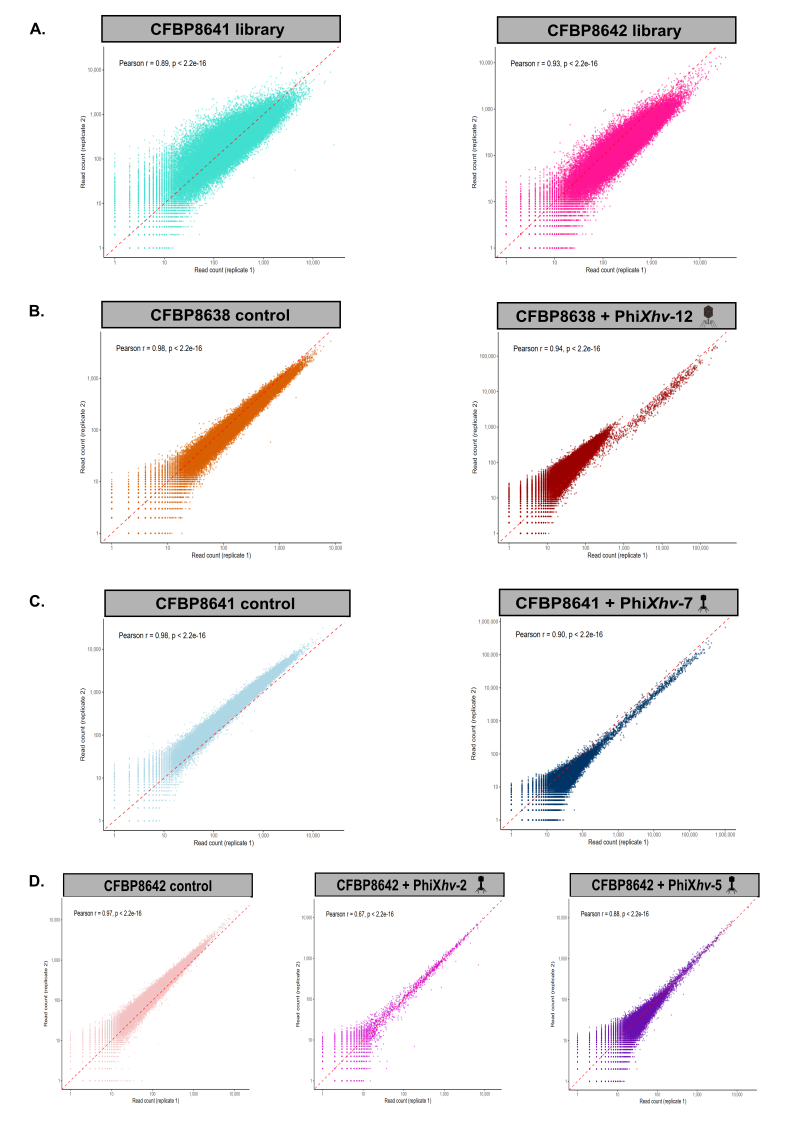

**Figure S4. Biological reproducibility of Tn-seq datasets.** Pairwise correlations of gene-level read counts between two biological replicates are shown for each experimental condition. Each dot represents a gene, with axes indicating the read count obtained for that gene in biological replicate 1 and biological replicate 2. Pearson correlation coefficients (*r*) and associated *p. values* are indicated in each plot. The red dashed line indicates the identity line (y=x). A. Correlation plots for the novel transposon mutant libraries of strains CFBP8641 and CFBP8642. **(B-D)** Correlation plots for untreated controls and corresponding phage-treated libraries. **B.** CFBP8638 control and CFBP8638 library following challenge with Phi*Xhv*-12. **C.** CFBP8641 control and CFBP8641 library following challenge with Phi*Xhv*-7. **D.** CFBP8642 control and CFBP8642 library following challenge with Phi*Xhv*-2 or Phi*Xhv*-5. Untreated controls were grown in TSB without phage.

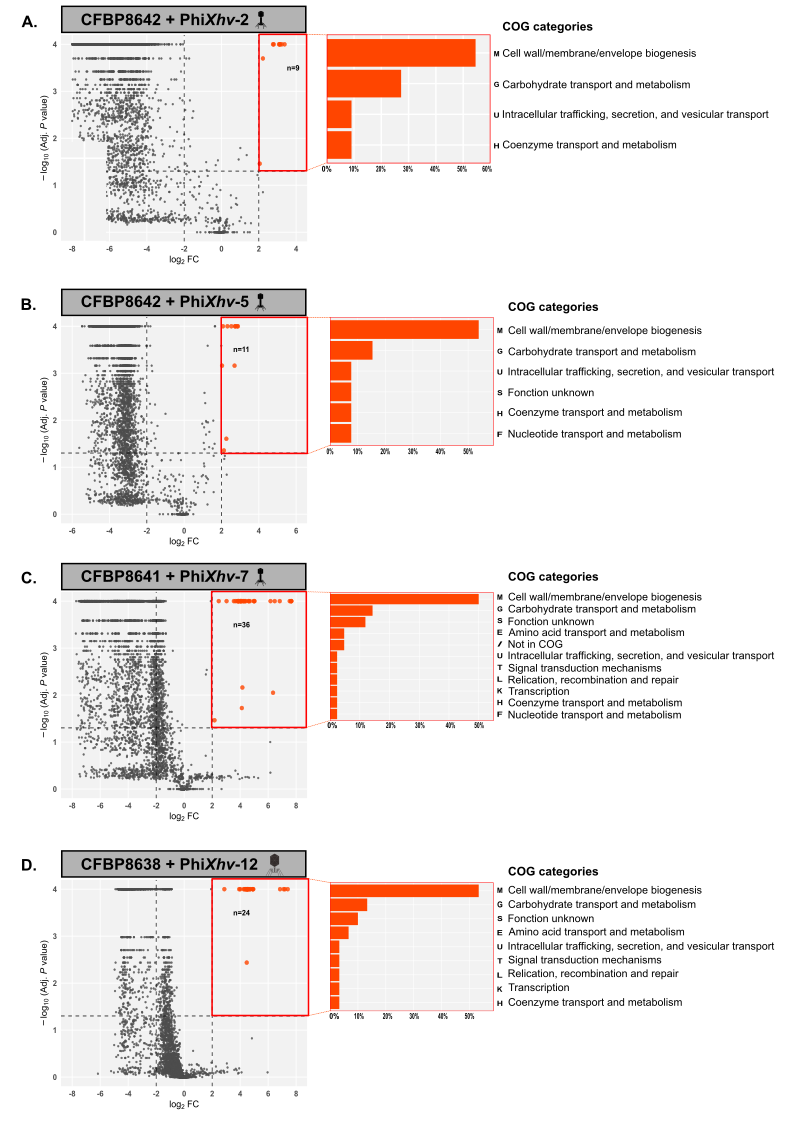

**Figure S5. Identification and functional classification of *X*. *hortorum* pv. *vitians* genes associated with successful infection by four phages of the cocktail using Tn-seq.** Volcano plots show the differential abundance of transposon mutants between untreated control and phage-treated libraries. Significantly enriched genes (log_2_FC > 2, *q*. *value* < 0.05) are highlighted in orange, and the number of significantly enriched genes (*n*) is indicated for each dataset. Dashed lines indicate the significance and log_2_FC thresholds used to identify enriched genes. Corresponding bar plots show the distribution of significantly enriched genes among functional categories based on Clusters of Orthologous Groups (COG) annotations. (A) Phi*Xhv*-2, screened against the CFBP8642 transposon library, (B) Phi*Xhv*-5, screened against the CFBP8642 transposon library, (C) Phi*Xhv*-7, screened against the CFBP8641 transposon library, (D) Phi*Xhv*-12, screened against the CFBP8638 transposon library.

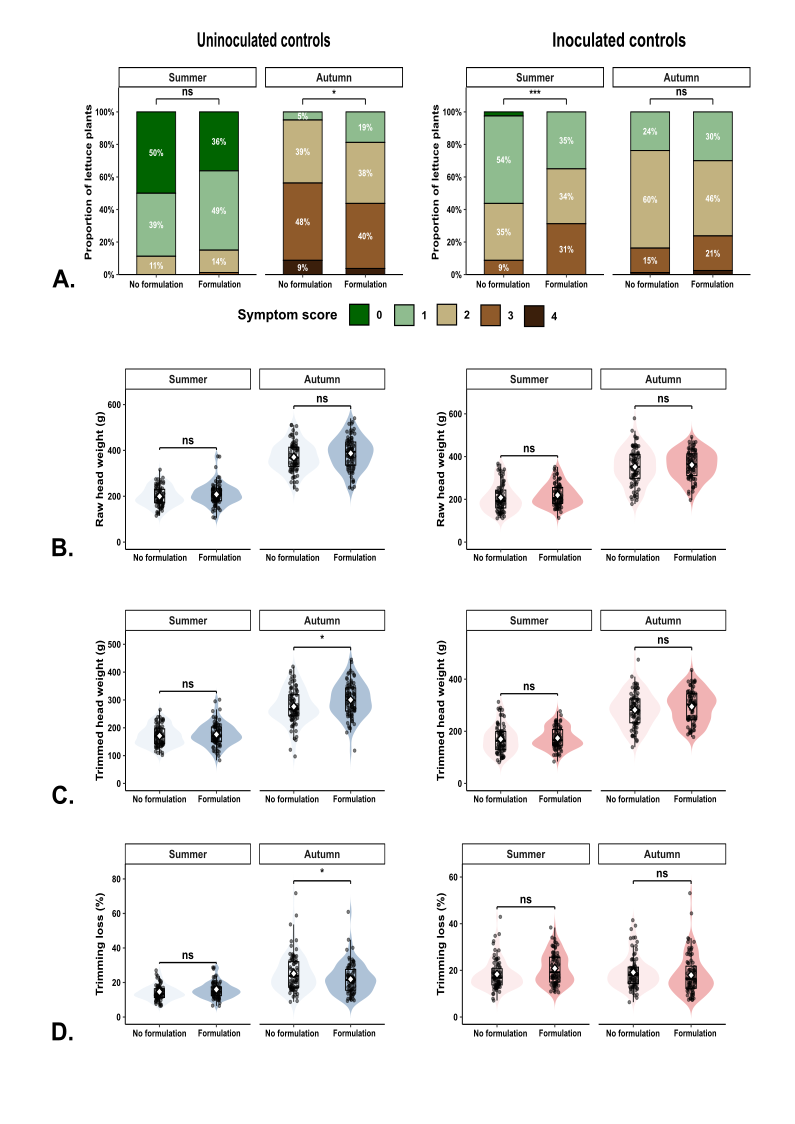

**Figure S6. Effect of the formulation on disease severity and yield-related traits in the field trials.** **A.** Distribution of disease severity scores at harvest in uninoculated and inoculated controls, with or without formulation, during the summer and autumn field trials. Stacked bars represent the proportion of lettuce plants in each disease severity class (0-4), with percentages indicated within each segment. **B.** Raw head weight at harvest. **C.** Trimmed head weight after removal of damaged outer leaves. **D.** Trimming loss, express as a percentage of raw head weight. For each parameter, formulation and no-formulation treatments are compared separately for uninoculated and inoculated controls in each season. Violin plots show the distribution of individual plant values (dots), with embedded boxplots indicating medians and interquartile ranges. Asterisks indicate significant differences between formulation and no-formulation treatments (*p* < 0.05 (*), *p* < 0.01 (**), *p* < 0.001, ns = not significant).
